# Growth factor-free synthetic matrices reveal intrinsic skeletal progenitor competence

**DOI:** 10.64898/2026.06.11.731042

**Authors:** Q Vallmajo-Martin, E Avilla-Royo, L Moser, D Alpern, V Gardeux, BM Carrara, A Barbero, I Martin, B Deplancke, M Lütolf, M Ehrbar

## Abstract

Clinical translation of human skeletal progenitor cells has been limited by the inability to reliably predict their regenerative performance, as current characterization methods rely on phenotypic markers or instructive differentiation assays that obscure intrinsic cell function. Here we demonstrate that human skeletal progenitor competence can only be revealed within a precisely defined and growth factor-free synthetic microenvironment. By independently tuning matrix mechanics, degradability, adhesion, and cell density within a fully synthetic poly(ethylene glycol) hydrogel, we create a permissive niche that enables human bone marrow stromal cells (hBMSCs) to undergo spontaneous bone formation *in vivo* without osteoinductive supplementation. These exogenous growth factor-free conditions expose donor-specific differences in bone forming capacity that are otherwise masked. Strikingly, supplementation with even low concentrations of BMP-2 abolished inter-donor variability, directly demonstrating that osteoinductive factors override intrinsic progenitor competence. Single-cell transcriptomics, proteomic profiling, and longitudinal *in vivo* analyses converge to show that high performing donors are defined by chondrocyte-primed transcriptional states and coordinated extracellular matrix (ECM) remodeling programs that establish a pro-osteochondral niche and drive bone formation via endochondral ossification. In contrast, low performing donors fail to initiate this cascade despite comparable phenotypic profiles and proliferation rates. Together, these findings implicate matrix context as a critical regulator of stem cell competence. Within this permissive synthetic niche, intrinsic ECM remodeling capacity emerges as a defining feature of donor-specific osteogenic potential, one that remains invisible to conventional assays. These matrices thus provide a functional platform for prospective stratification of skeletal progenitor cells, with direct implications for donor selection in regenerative medicine.

**Graphical Abstract:** 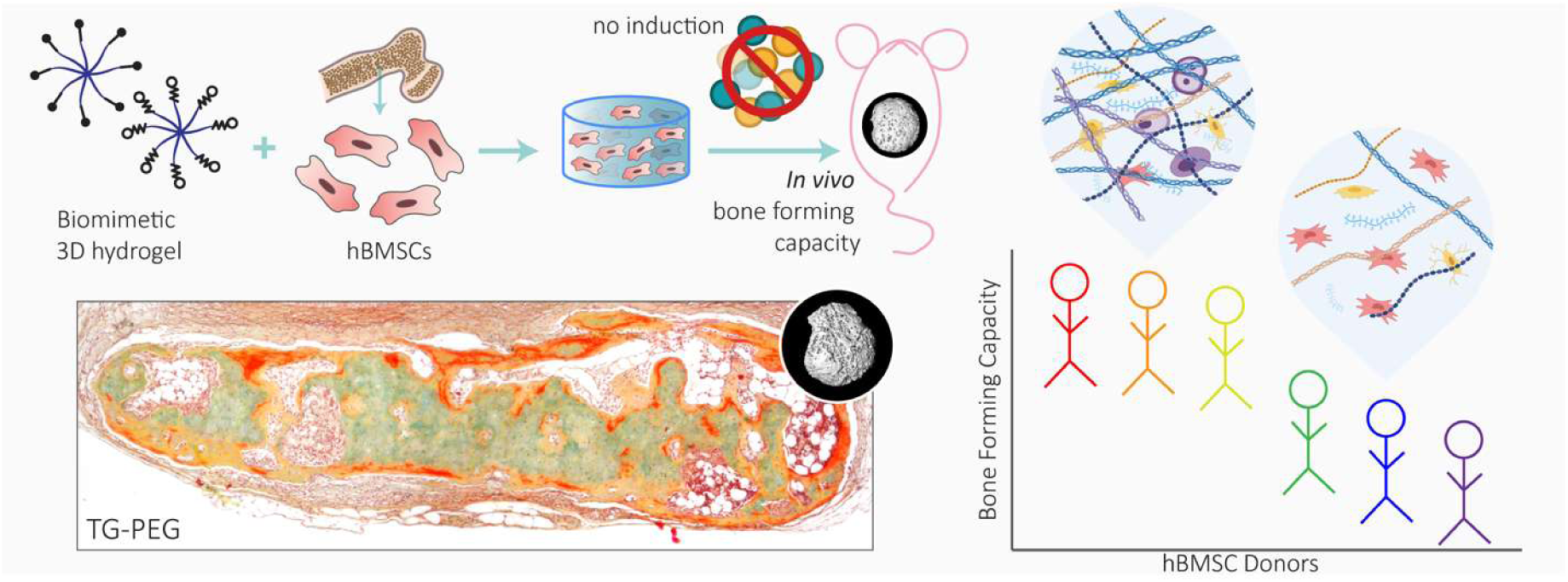

## Main

Skeletal stem and progenitor cells have long been recognized as central regulators of bone formation and repair, yet their precise identity and function remain unresolved^1–4^. Human bone marrow stromal cells (hBMSCs) have been widely employed in regenerative medicine due to their capacity to support bone formation in preclinical models^5,6^. However, their clinical translation has produced inconsistent outcomes, largely owing to the lack of specific and reliable surface markers that can distinguish *bona fide* skeletal progenitor subpopulations^7–10^. Consequently, hBMSC capacity is typically inferred from *in vitro* differentiation assays that poorly mimic physiological or therapeutic contexts^11^. What is lacking is a system that interrogates hBMSC function without imposing its own biological agenda, in other words, one that permits rather than prescribes differentiation.

Ideally, the functional potential of skeletal stem and progenitor cells would be evaluated within a defined microenvironment that permits controlled, reciprocal interactions between cells and their surrounding matrix^12–15^. Numerous biomaterials have been explored for this purpose, including hydroxyapatite/β-tricalcium phosphate (HA/TCP) ceramics scaffolds^16,17^ and naturally derived matrices such as collagen^18,19^, fibrin^20^, or extracellular matrix (ECM)-based hydrogels^1,21,22^. Yet these biomaterials bear inherent osteoconductive or bioactive properties that exogenously instruct rather than reveal cell behavior, thereby masking the intrinsic differentiation programs of the encapsulated cells^12,13^.

To address these gaps, we established a minimalistic *in vivo* system, free of exogenous osteoinductive factors, to directly assess the intrinsic bone-forming capacity of hBMSCs. Building on a modular, chemically defined poly(ethylene glycol) (PEG)-based hydrogel^23–28^, we systematically tuned matrix stiffness, degradability, and cell-adhesion properties to create a neutral yet permissive environment that supports spontaneous bone and bone marrow formation without supplementation of any growth factors such as bone morphogenetic protein 2 (BMP-2). This fully synthetic platform directly interrogates human skeletal progenitor function *in vivo*, revealing that bone formation arises via an intrinsic endochondral ossification program. Comparative analyses across donors further uncover that, within this synthetic context, chondrogenic ECM deposition capacity distinguishes high- from low-performing hBMSCs, highlighting how cell-intrinsic differentiation potential can be exposed within a fully defined synthetic environment.

### Rational design of a minimal synthetic environment to test intrinsic bone-forming capacity of hBMSCs

The intrinsic regenerative potential of skeletal progenitors can only be revealed under *in vivo* environments free of exogenous instructive cues. For this purpose, we engineered a chemically defined poly(ethylene glycol)-based hydrogel that is crosslinked by transglutaminase FXIII (referred to as TG-PEG^23^). This hydrogel platform permits independent tuning of stiffness, proteolytic degradability, and cell adhesion (**Fig. 1a**). First, we compared the ability of TG-PEG to promote hBMSC-mediated formation of ectopic bone ossicles in mice with naturally-derived collagen, fibrin, and ECM-based (named as ECMatrix) hydrogels. hBMSCs (at a concentration of 20x10^6^ cells per ml of hydrogel) were encapsulated in TG-PEG or in natural matrices with matched stiffness equivalent to a storage modulus of ca. 50 Pa (**Fig. 1b** *in grey*) and immediately implanted subcutaneously in mice. Micro-computed tomography (microCT) evaluations 8 weeks post-implantation showed that collagen scaffolds failed to form bone, while small bony ossicles were formed in fibrin (0.187 ± 0.016 mm^3^), ECMatrix (0.442 ± 0.467 mm^3^), and low stiffness TG-PEG (0.313 ± 0.129 mm^3^) matrices (**Fig. 1c** *in grey*). Modified Movat’s pentachrome (MP) stained histological sections confirmed the formation of an ECM rich in glycosaminoglycans (GAGs) and bone tissues within these soft scaffolds (**Fig. 1d**). However, all hydrogels revealed an 80 to 100% reduction in size, suggesting the resorption of these scaffolds driven by cell proteolytic activity, consistent with reports in fibrin hydrogels^29^.

**Figure 1.**
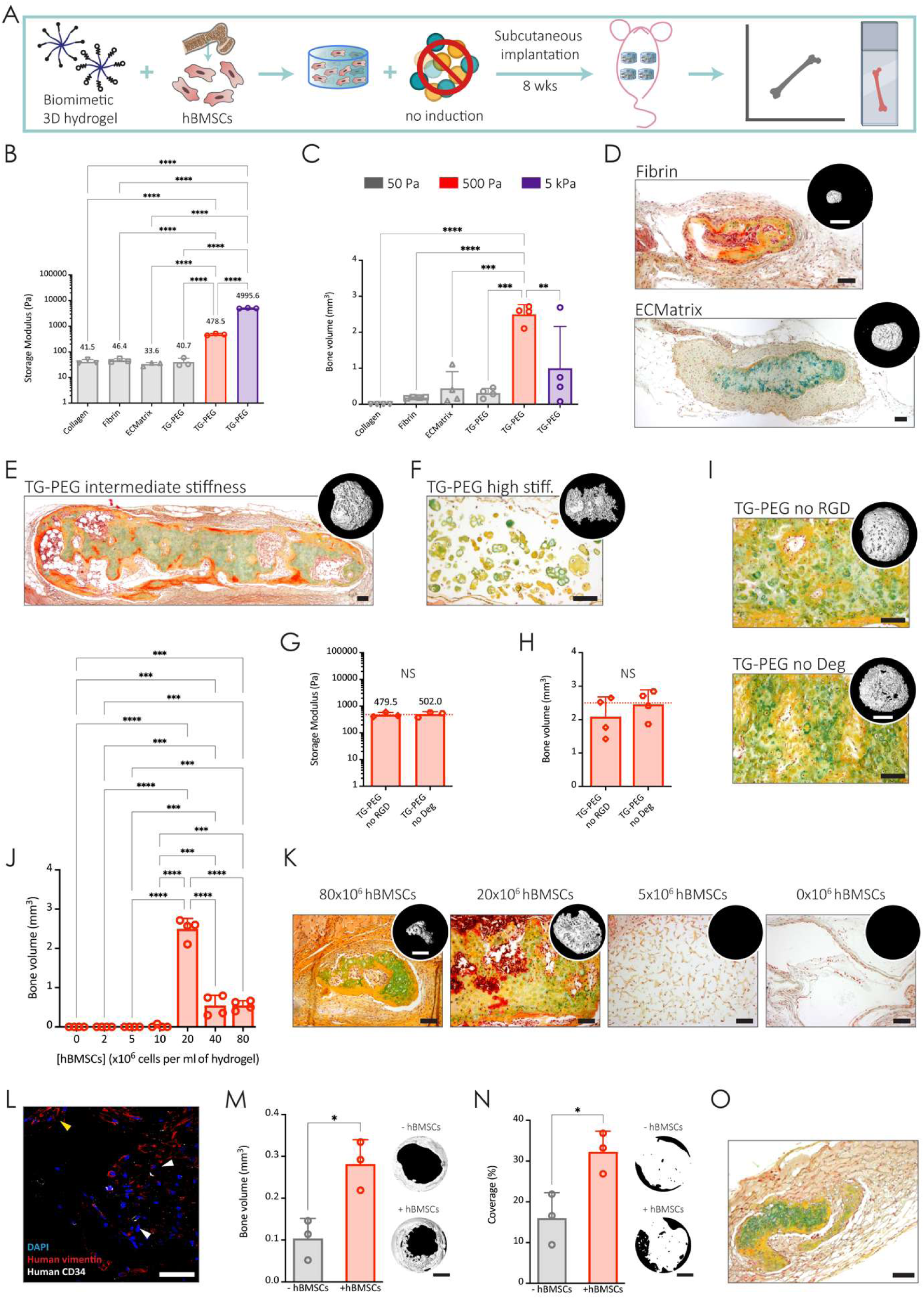
A minimally instructive synthetic matrix enables growth-factor-free bone ossicle and marrow niche formation. **a,** Schematic of the experimental workflow. Human bone marrow stromal cells (hBMSCs) were encapsulated in three-dimensional (3D) growth-factor-free biomimetic hydrogels and implanted subcutaneously in immunodeficient mice. After 8 weeks, explanted ossicles were analyzed by micro-computed tomography (microCT) and histological assessment. Created with BioRender.com. b, Storage moduli of collagen, fibrin, ECMatrix, and TG-PEG hydrogels of low (50 Pa), intermediate (500 Pa), or high stiffness (5 kPa), measured by rheometry (n = 3 independent hydrogel measurements per condition). Mean values are indicated above bars. c, Quantification of total bone volume measured by microCT in ossicles formed from different 3D matrices containing hBMSCs (20 x 10⁶ cells ml⁻¹) at 8 weeks post-implantation (n = 4 implants per condition). d-f, Representative modified Movat’s pentachrome (MP)-stained sections of ossicles formed in low-stiffness matrices (**d**), intermediate-stiffness TG-PEG hydrogels (**e**), and high-stiffness TG-PEG hydrogels (**f**). Insets show corresponding microCT reconstructions. Scale bars: 100 µm (MP histology); 2 mm (microCT). **g-i,** Effect of cell-matrix interactions on ossicle formation. hBMSCs (20 x 10⁶ cells ml⁻¹) were encapsulated in intermediate-stiffness TG-PEG hydrogels lacking RGD adhesion motifs or matrix metalloproteinase (MMP)-sensitive degradable sites. **g,** Storage moduli measured by rheometry (n = 3 independent hydrogel measurements per condition). Mean values are indicated above bars. Dotted line in red indicates the average storage moduli of intermediate-stiffness TG-PEG hydrogels with RGD adhesion motifs and MMP-sensitive sites. **h,** Bone volume quantified by microCT at 8 weeks post-implantation (n = 4 implants per condition). Dotted line in red indicates the average bone volume of intermediate-stiffness TG-PEG hydrogels with RGD adhesion motifs and MMP-sensitive sites. **i,** Representative MP-stained sections with corresponding microCT reconstructions. Scale bars: 100 µm (MP histology); 2 mm (microCT). **j-k,** Effect of cell seeding density on ossicle formation. hBMSCs were encapsulated at varying concentrations in intermediate-stiffness TG-PEG hydrogels containing RGD and MMP-sensitive sites. **j,** Bone volume quantified by microCT at 8 weeks (n = 4 implants per condition). **k,** Representative MP-stained sections with corresponding microCT reconstructions. Scale bars: 100 µm (MP histology); 2 mm (microCT). **l,** Functional bone marrow formation within engineered ossicles. Following ossicle formation in NSG mice, human hematopoietic stem and progenitor cells (hHSPCs) were systemically injected via the tail vein. Humanization was observed 16 weeks post-transplantation, with human vimentin⁺ hBMSCs localized to bony regions (yellow arrow) and human CD34⁺ hHSPCs residing within the marrow cavity (white arrows). Nuclei were counterstained with DAPI, human-specific vimentin in red, and human-specific CD34 in white. Scale bar: 50 µm. **m-o,** Regeneration of critical-size calvarial defects using growth-factor-free TG-PEG hydrogels. Optimized intermediate-stiffness TG-PEG hydrogels containing RGD and MMP-sensitive sites were implanted with or without hBMSCs (20 x 10⁶ cells ml⁻¹) into murine calvarial defects for 4 weeks. **m,** Bone volume quantified by microCT (n = 3 implants per condition). Representative reconstructions are shown. Scale bars: 2 mm. **n,** Defect coverage quantified by microCT (n = 3 implants per condition). Representative reconstructions are shown. Scale bars: 2 mm. **o,** Representative MP-stained section confirming newly formed bone tissue within the defect area in the calvaria bone. Scale bar: 100 µm, image shows all defect area. In MP staining, glycosaminoglycan-rich cartilage appears blue/green, collagenous bone matrix appears yellow, vascularized tissue and marrow appear red, and nuclei appear black. Bars represent mean ± s.d.; dots indicate individual implanted hydrogels. Unpaired two-tailed Student’s t-test (g, h, m, n); One-way ANOVA with Tukey’s multiple comparisons test (b, c, j). * P < 0.05; ** P < 0.01; *** P < 0.001; **** P < 0.0001; NS, not significant.

### A minimally instructive synthetic matrix enables growth-factor-free bone ossicle formation

Next, we hypothesized that hydrogels with higher crosslinking density and stiffness would better promote ossicle development by providing more stability to the delivered cells. In contrast to natural hydrogels and owing to their synthetic nature, TG-PEG hydrogels are easily customizable over a wide range (100-fold) of stiffnesses by increasing the initial concentration of the polymer backbone^30^ (**Fig. 1b** *in red*). Strikingly, TG-PEG hydrogels of intermediate stiffness (storage modulus of 500 Pa) showed significantly increased bone formation (2.089 ± 0.591 mm^3^) without any osteogenic supplements (**Fig. 1c** *in red*). MicroCT evaluations confirmed the *de novo* formation of mineralized tissue, while MP staining revealed the coexistence of bone, cartilage, and marrow-like regions, indicating that the synthetic hydrogel of intermediate stiffness permitted bone formation in the absence of inductive factors (**Fig. 1e**). In the stiffest hydrogels tested (storage modulus of 5 kPa) (**Fig. 1b** *in purple*), however, the capability of hBMSCs to remodel the hydrogel and deposit their own mineralized ECM was compromised, resulting in highly variable bone volumes (1.000 ± 1.160 mm^3^) (**Fig. 1c** *in purple*). Histological analysis by MP staining indicated that transplanted cells in stiffest hydrogels acquired a cartilage-like morphology and highlighted the lack of diverse cell types in the constructs, indicative of low host cell infiltration (**Fig. 1f**). Together, these findings emphasize the need to precisely tune hydrogel stiffness to promote osteogenesis and reveal that an intermediate stiffness of 500 Pa most effectively supports bone formation within our system.

Having established that intermediate stiffness is required for productive ossicle formation, we next asked which biochemical cues within this mechanical window are necessary for bone niche establishment. Cells establish and remodel their microenvironment through iterative, reciprocal crosstalk that involves ECM degradation via secretion of matrix metalloproteinases (MMPs)^31^ and deposition of newly synthesized ECM components^32^. These deposited ECM proteins contain cell-adhesion peptides, such as RGD, that enable cell adhesion and spreading^33^. Therefore, to dissect the role of cell-matrix interactions in this setup, we engineered TG-PEG hydrogels of intermediate stiffness that either lacked RGD sites or the MMP-sensitive backbone. We first confirmed that engineering the hydrogel backbone did not change the overall stiffness (**Fig. 1g**). Surprisingly, total bone volume was not affected by omitting RGD binding sites or MMP-degradability, accomplishing 2.089 ± 0.591 and 2.456 ± 0.433 mm^3^, respectively (**Fig. 1h**). However, histological evaluations revealed that in the absence of RGD or MMP-degradable motifs, hBMSCs deposited an ECM rich in GAGs and adopted a round, chondrocyte-like phenotype similar to that observed in high stiffness hydrogels. Additionally, under these conditions, the ossicles lacked a bone marrow cavity containing host immune cells (**Fig. 1i**). This was in stark contrast to the robust marrow establishment observed in the TG-PEG formulations containing RGD and MMP-degradation sites (**Fig. 1e**). These findings are consistent with prior studies showing that integrin-mediated adhesion^34^ and matrix remodeling^30^ support osteogenic tissue formation, and further demonstrate in our fully synthetic system that matrix adhesiveness and degradability are both required to enable host cell infiltration and subsequent bone marrow niche formation. Collectively, these results demonstrate that independent tuning of stiffness, MMP-sensitivity, and cell adhesion, an approach only achievable in synthetic but not in natural biomaterials, is critical for the evaluation of hBMSC bone forming capacity.

Next, we tested whether our *in vivo* engineered ossicles could be further optimized by varying the number of encapsulated hBMSCs in optimized TG-PEG hydrogels (i.e. storage modulus of 500 Pa, 50 µM RGD, and containing the MMP-1-sensitive degradable sequence). While by increasing the seeding density (40 and 80x10^6^ cells/ml) the amount of formed bone significantly decreased, by decreasing the seeding density (10x10^6^ cells/ml and lower) bone formation was no longer obtained (**Fig. 1j**). MP stained tissue sections of implants seeded with the highest number of hBMSCs (80x10^6^ cells/ml) revealed small bone ossicles that were filled with densely packed chondrocytes (**Fig. 1k**). Whereas in lower seeding densities (5x10^6^ cells/ml) loose cellular networks were observed, even after 8 weeks of *in vivo* implantation. This indicates the need for balanced levels of hBMSC-driven tissue formation and hydrogel degradation^35^, which is mediated by their secreted soluble factors and deposited ECM components.

These results identify an optimal formulation in which the TG-PEG hydrogel enables tissue morphogenesis while remaining permissive to host infiltration and cellular remodeling without need of any bone-inducing supplementation such as BMP-2. Together, these results establish that matrix permissivity, defined by the precise balance of mechanical support, proteolytic accessibility, and adhesive signaling, is the critical materials parameter enabling cell-intrinsic bone formation, a design principle inaccessible to naturally derived matrices.

### Synthetic ossicles support functional hematopoietic niches and repair critical bone defects

To investigate the functionality of the bone marrow compartment^36^, we induced bone ossicles in NSG mice 8 weeks before we systemically injected human hematopoietic stem and progenitor cells (hHSPCs) (**Supplementary Fig. 1a**). Histological analysis 16 weeks post-transplantation revealed that bone ossicles comprised a marrow containing engrafted human specific CD34^+^ HSPCs (**Fig. 1l**), indicating the functionality of the engineered bone-marrow niches.

To test whether intrinsic hBMSC bone-forming capacity is sufficient to repair critical bone defects, we implanted hBMSC-laden TG-PEG hydrogels into critical-sized calvarial defects^28,37,38^ in the absence of BMP-2 (**Supplementary Fig. 1b**). This resulted in a significant 2.7-fold increase in bone formation and defect coverage compared to acellular controls (**Fig. 1m, n**), with histological analysis confirming direct participation of implanted hBMSCs in new bone deposition (**Fig. 1o**). Notably, this was achieved without any osteoinductive supplementation, a clinically significant finding given the well-documented adverse effects of BMP-2 in craniofacial applications, including ectopic bone formation, inflammation, and nerve compression^39^. These results demonstrate that the intrinsic bone-forming capacity of hBMSCs is sufficient to drive repair of bone defects.

### Bone ossicle formation is cell-autonomous and paradoxically suppressed by exogenous osteoinductive priming

We next examined whether the transplanted hBMSCs or the recruited host cells were responsible for directing bone ossicle formation. To do so, we engineered the optimized TG-PEG hydrogels (i.e. storage modulus of 500 Pa, 50 µM RGD, and containing the MMP-1-sensitive degradable sequence) to locally present in their microenvironment cell-surface ligands known to inhibit hBMSC differentiation. Jagged1 or Delta-like-4 (DLL4), two Notch signaling ligands known to maintain the pool of mesenchymal progenitors in the bone marrow^40^, were modularly incorporated into the hydrogel backbone using a high affinity protein A-based binding strategy^27,41^ (**Fig. 2a**). Jagged1 or DLL4 presenting TG-PEG hydrogels completely abolished bone formation (**Fig. 2b** and **2c**), directly demonstrating that ossicle development depends on hBMSC differentiation rather than recruitment and activation of host progenitors. This confirmed that the bone-forming capacity observed in this system is cell-autonomous, intrinsic to the transplanted hBMSCs, and not a consequence of scaffold-mediated host cell recruitment.

**Figure 2.**
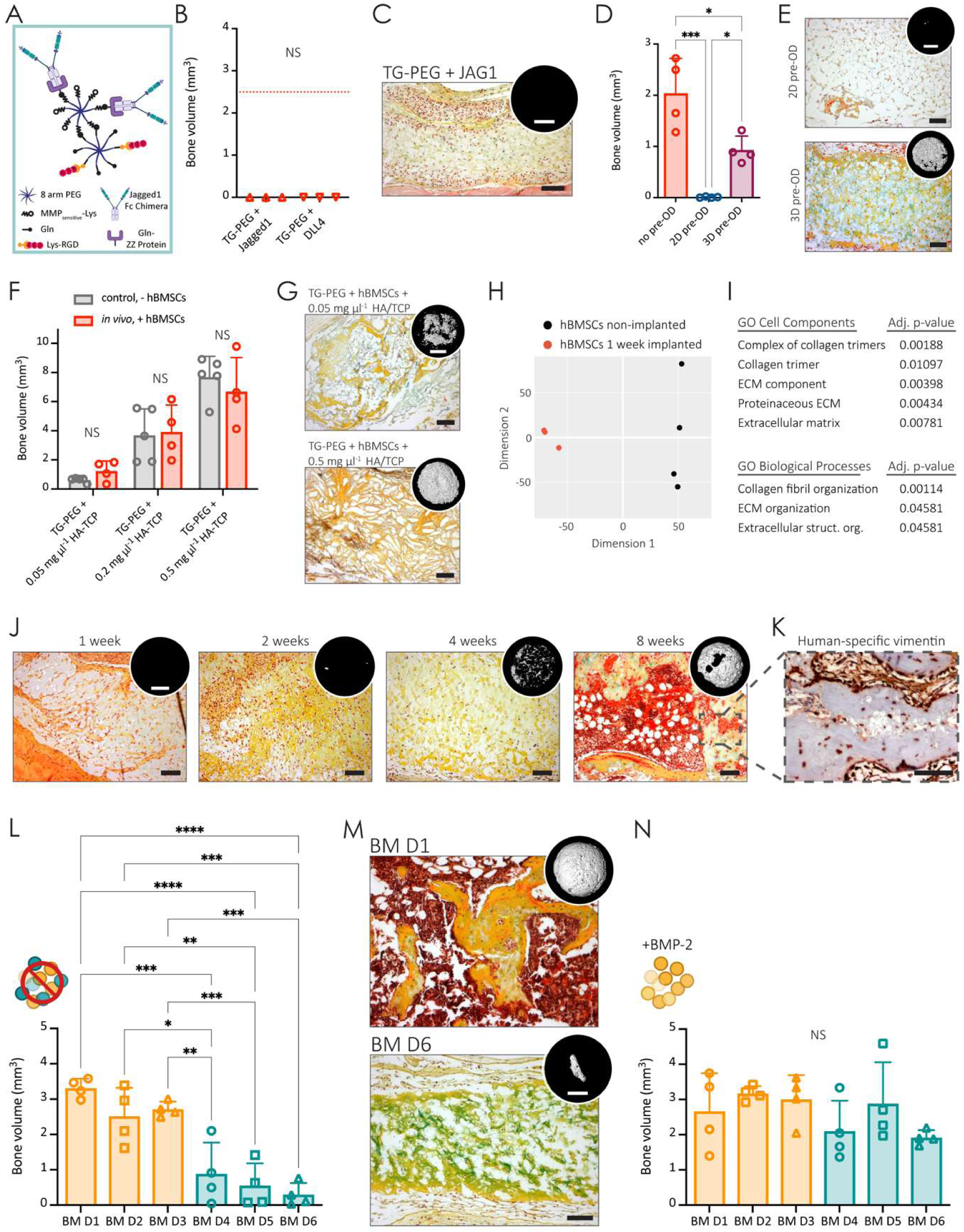
A growth-factor-free synthetic environment exposes cell-autonomous bone formation and intrinsic donor variability. **a-c,** Effect of Notch ligands presentation on hBMSC-mediated ossicle formation. hBMSCs (20 x 10⁶ cells ml⁻¹) were encapsulated in intermediate-stiffness TG-PEG hydrogels containing RGD adhesion motifs and MMP-sensitive degradable sites. a, Schematic illustrating modular TG-PEG hydrogel functionalization for immobilization of Notch ligands in addition to MMP-sensitive sites and RGD integrin-binding motifs. Recombinant Jagged1-Fc or DLL4-Fc proteins were covalently tethered to the TG-PEG backbone using a protein A-based Gln-ZZ linker strategy, as previously described^27,41^. The ZZ domain binds Fc-tagged proteins with high affinity, whereas the Gln domain enables covalent incorporation into the TG-PEG hydrogel during crosslinking with the transglutaminase Factor XIIIa. Created with BioRender.com. b, Bone volume quantified by microCT at 8 weeks post-implantation (n = 3 implants per condition). Dotted line in red indicates the average bone volume of intermediate-stiffness TG-PEG hydrogels with RGD adhesion motifs and MMP-sensitive sites. c, Representative modified MP-stained sections with corresponding microCT reconstructions. Scale bars: 100 µm (MP histology); 2 mm (microCT). d-e, Effect of osteogenic pre-differentiation on ossicle formation. hBMSCs were cultured for 8 days in osteogenic medium either in 2D culture or within 3D TG-PEG hydrogels prior to subcutaneous implantation. Pre-differentiated 2D cultures were re-encapsulated before implantation, whereas 3D cultures were implanted directly. d, Bone volume quantified by microCT after 8 weeks (n = 4 implants per condition). e, Representative MP-stained sections with corresponding microCT reconstructions. Scale bars: 100 µm (MP histology); 2 mm (microCT). f-g, Effect of osteoconductive particle incorporation on bone formation. Increasing amounts of osteoconductive hydroxyapatite/β-tricalcium phosphate (HA/TCP; 60/40, particle diameter <75 µm) were incorporated into intermediate-stiffness TG-PEG hydrogels containing hBMSCs prior to implantation. f, Bone volume was quantified by microCT at 8 weeks (red, n = 4 implants per condition) and compared to non-implanted hydrogel controls containing only particles (grey, n = 5 hydrogels per condition). g, Representative MP-stained sections with corresponding microCT reconstructions. Scale bars: 100 µm (MP histology); 2 mm (microCT). **h-i,** Early transcriptional response of hBMSCs to *in vivo* implantation. Human cells were isolated from TG-PEG hydrogels 1 week post-implantation (n = 3 implants) and compared to controls of non-implanted hBMSCs in hydrogels (n = 4 hydrogels) by bulk RNA sequencing. **h,** Principal component analysis (PCA). **i,** Gene ontology enrichment analysis of differentially expressed genes, showing cellular components and biological processes. **j,** Longitudinal assessment of hBMSC-mediated ossicle development. Representative MP-stained sections at indicated time points with corresponding microCT reconstructions. Scale bars: 100 µm (MP histology); 2 mm (microCT). **k,** Representative immunohistochemical staining for human-specific vimentin identifying human cells within an ossicle after 8 weeks of implantation. Scale bar: 100 µm. **l,** Donor-dependent differences in bone-forming capacity revealed by the growth-factor-free TG-PEG system. Bone volume quantified by microCT at 8 weeks post-implantation in hBMSCs derived from six healthy bone marrow donors (BM D1-BM D6; N = 6 donors; n = 4 implants per donor). **m,** Representative histological comparison of high- (BM D1) and low-performing (BM D6) donors. MP-stained sections illustrating differences in tissue remodeling (scale bar: 100 µm). Upper insets are the corresponding microCT reconstructions (scale bar: 2 mm). **n,** Effect of BMP-2 supplementation on donor-dependent variability. Bone volume quantified by microCT after implantation of hBMSCs in the presence of low-dose BMP-2 (12.5 ng µl⁻¹). BMP-2 supplementation abolished statistically significant inter-donor differences in bone volume (p = 0.7735 for BM D1 vs BM D6), demonstrating that osteoinductive growth factors override intrinsic donor-specific bone-forming capacity (N = 6 donors; n = 4 implants per donor). In MP staining, glycosaminoglycan-rich cartilage appears blue/green, collagenous bone matrix appears yellow, vascularized tissue and marrow appear red, and nuclei appear black. Bars represent mean ± s.d.; dots indicate individual implanted hydrogels. Unpaired two-tailed Student’s t-test (b); One-way ANOVA with Tukey’s multiple comparisons test (d, l, n). Two-way ANOVA with Šídák’s multiple comparisons test (f). * P < 0.05; ** P < 0.01; *** P < 0.001; **** P < 0.0001; NS, not significant.

To test whether inducing hBMSC condensation or osteogenic differentiation prior to implantation^42^ could further improve ossicle formation, we pre-differentiated hBMSCs either in 2D or already in 3D TG-PEG hydrogels for 8 days prior to *in vivo* implantation (**Supplementary Fig. 2a**). Surprisingly, in both cases, pre-differentiated hBMSCs resulted in significantly reduced bone formation compared to undifferentiated hBMSCs (**Fig. 2d** and **2e**). We also tested whether the addition of osteoconductive hydroxyapatite tricalcium phosphate particles (HA/TCP, 60/40, diameter < 75 µm), commonly used to direct osteogenic differentiation *in vivo* and in the clinical setting^43^, into the TG-PEG hydrogels would augment bone ossicle formation by hBMSCs (**Supplementary Fig. 2b**). However, despite the stiffening effect particles had on the TG-PEG constructs (**Supplementary Fig. 2c, d**), hBMSCs were not able to remodel the implants into bone ossicles (**Fig. 2f** and **2g**). These results reinforced that the herein presented TG-PEG hydrogel formulation is sufficient to allow hBMSCs to unfold their inherent bone formation capacity, and even more that osteoinductive or osteoconductive supplementation in these optimized conditions compromise hBMSC osteogenic remodeling.

To resolve the molecular events accompanying differentiation of hBMSCs encapsulated in TG-PEG *in vivo*, we performed transcription profiling of hBMSCs at two developmental stages, prior to and one week after subcutaneous implantation. Bulk RNA barcoding and sequencing (BRB-seq)^44^ analysis displayed profound differences in hBMSCs after just one week post-implantation, as shown by separate clustering in principal component analysis (PCA, **Fig. 2h**). More specifically, GO enrichment and individual gene analysis revealed rapid upregulation of a coordinated cartilage-matrix program within just one week of implantation (**Fig. 2i** and **Supplementary Fig. 2e, f**). This included fibrillar collagens characteristic of hyaline cartilage (collagen type II alpha 1 chain, COL2A1 and collagen type XI alpha 2 chain, COL11A2), the large aggregating proteoglycan aggrecan (ACAN) that provides compressive resistance to developing cartilage, the matrix organizer hyaluronan and proteoglycan link protein 1 (HAPLN1) that stabilizes aggrecan-hyaluronan complexes in the pericellular niche, the small leucine-rich proteoglycan lumican (LUM) implicated in collagen fibrillogenesis, and the matrix remodeling-associated protein 5 (MXRA5)^45^. The coordinated induction of these genes points to spontaneous entry into a chondrogenic program, the molecular hallmark of endochondral ossification, without any exogenous inductive signal.

Longitudinal MP staining corroborated the rapid molecular changes, showing early ECM deposition in these forming ossicles (**Fig. 2j**). Already after 1 week of implantation, hydrogels were rich in GAGs that from week 4 on remodeled into calcified collagen fibers as seen by microCT reconstructions. Up to eight weeks after implantation, human cells were found in the ossicles in endosteal and stromal niches supporting the bone marrow (**Fig. 2k**). Previously, hBMSCs were shown to undergo endochondral ossification only after *in vitro* pre-differentiation with TGF-β3^18,19^ or systemic supplementation with daily doses of human parathyroid hormone^21^. Remarkably, here, we provide the first evidence that hBMSCs in the absence of inductive microenvironmental cues undergo bone formation via *in situ* endochondral ossification. Altogether, these results indicated that forcing hBMSCs to undergo directly osteogenic differentiation by *in vitro* pre-differentiation or osteoconductive particles may suppress *in vivo* endochondral ossification and therefore result in less efficient bone formation. This endochondral program operates without exogenous induction, but does it operate equally across donors?

### A growth-factor-free synthetic environment exposes intrinsic donor variability in bone-forming potential

To determine whether the TG-PEG platform captures donor-to-donor variation in bone-forming capacity, we measured the bone volume formed by hBMSCs from six healthy donors (females or males from 17 to 39 years old, **Supplementary Fig. 2g**). We found significant differences in bone volumes ranging from 3.313 ± 0.265 mm^3^ (BM D1) to 0.298 ± 0.327 mm^3^ (BM D6) uncorrelated to age or sex (**Fig. 2l**). Furthermore, histological analysis at 8 weeks of implantation confirmed that samples corresponding to higher bone volumes exhibited collagen-rich and fully developed bone ossicles containing an extensive bone marrow compartment, consistent with prior studies linking marrow cavity formation to haematopoietic niche establishment^46^. Implants with lower bone volumes showed no bone marrow formation (**Fig. 2m**). Strikingly, when encapsulating hBMSCs in presence of even low concentrations (12.5 ng µl⁻¹) of BMP-2, differences of formed bone volumes between donors were reduced and no longer significant (**Fig. 2n**; p-value BM D1 and BM D6 = 0.453), demonstrating that growth factor induction obscures the intrinsic potential of the encapsulated cells. Therefore, donor-dependent differences in bone-forming capacity within this system were detectable precisely because the TG-PEG platform does not rely on osteoconductive or osteoinductive factors that would otherwise override cell-intrinsic behavior.

### Single-cell transcriptomic profiling reveals chondrocyte-primed hBMSC subpopulations underlying donor-specific osteogenic potential

To resolve the cellular heterogeneity of hBMSCs across donors and to identify subpopulations that may have distinct bone-forming capacities, we performed single-cell RNA sequencing (scRNA-seq) on the different BM donors, building on prior single-cell atlases that have documented transcriptional diversity within hBMSC populations^3,47^. After quality control, 54,035 annotated cells were visualized using uniform manifold approximation and projection (UMAP), with each donor color-coded (**Fig. 3a**). To minimize technical variability and potential batch effects, we utilized Parse Biosciences’ combinatorial barcoding technology (Parse-seq), which enabled the simultaneous processing of all donor samples within combined sublibraries. By pooling samples early in the workflow, technical noise was uniformly distributed, ensuring that observed transcriptomic differences reflect intrinsic donor biology rather than experimental artifacts. Consequently, we intentionally refrained from applying computational batch normalization across donors, as these donor-specific variations represent the primary biological signal of interest.

**Figure 3.**
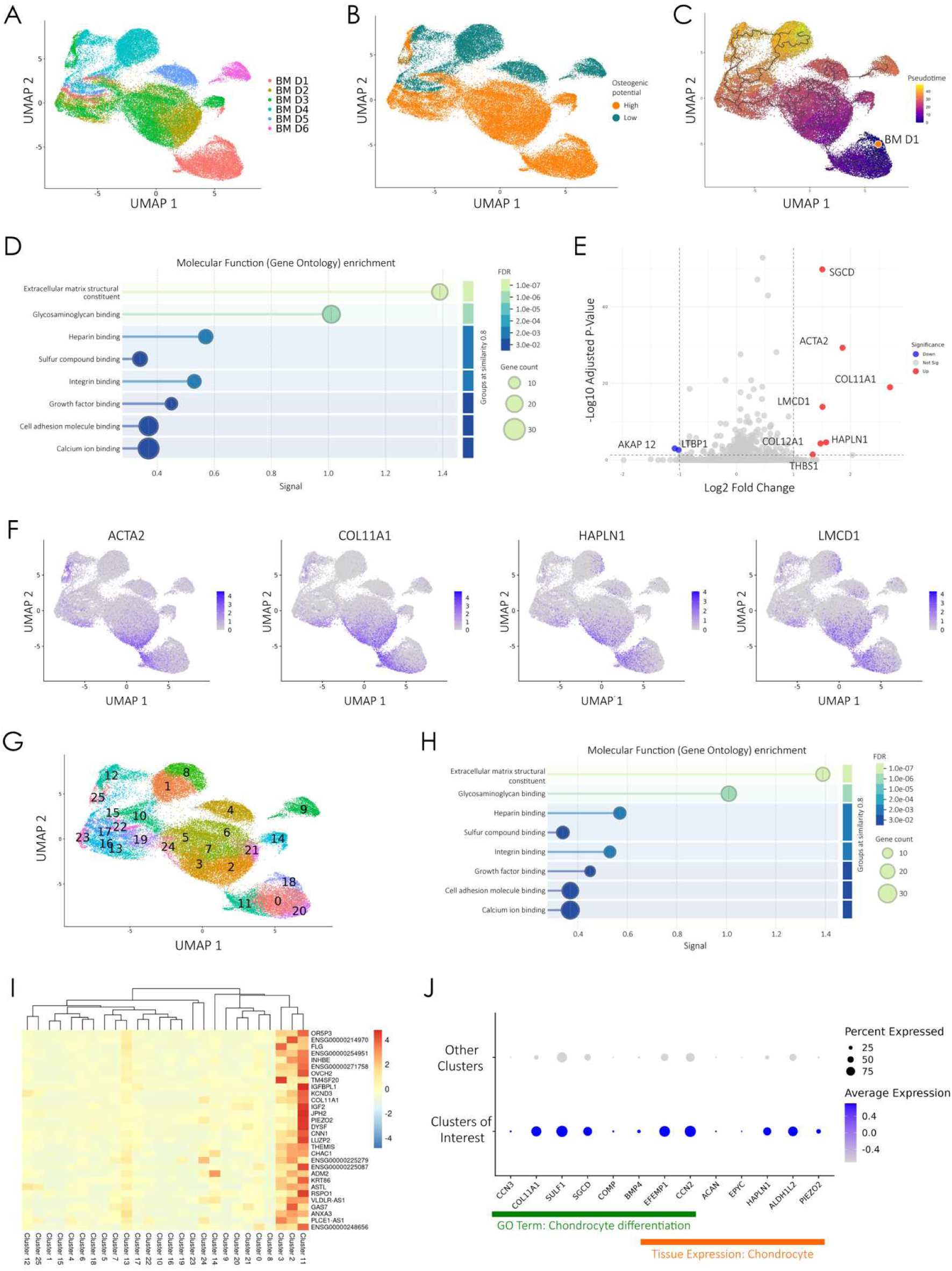
Single-cell transcriptomic profiling reveals hBMSC subpopulations underlying donor-specific osteogenic potential. **a,** Uniform manifold approximation and projection (UMAP) of all single cells after quality control, colored by donor identity. **b,** UMAP of the same dataset colored by osteogenic potential, with donors BM D1-D3 classified as high- and donors BM D4-D6 as low-osteogenic potential donors. **c,** Trajectory inference across all six donors with cells ordered along pseudotime. Root node was defined using BM D1 as the highest-performing donor. Color indicates pseudotime progression; point denotes the defined root of the trajectory, which is BM D1. **d,** Gene Ontology (GO) enrichment analysis for molecular function based on differentially expressed genes between high- and low-osteogenic-potential donors. Dot size indicates the number of genes associated with each term; color indicates false discovery rate (FDR). **e,** Volcano plot of pseudobulk differential gene expression comparing high- versus low-osteogenic-potential donors. Significantly regulated genes are highlighted (adjusted p < 0.05 and |log₂ fold change| > 1). **f,** UMAP feature plots showing expression of selected genes (*ACTA2*, *COL11A1*, *HAPLN1*, and *LMCD1*) across all cells. **g,** Single-cell clustering using shared nearest neighbor (SNN) modularity, identifying 25 transcriptionally distinct clusters colored individually. **h,** GO enrichment analysis for molecular function comparing combined clusters 2, 3, and 11 to all remaining clusters. Dot size reflects gene count; color indicates FDR. **i,** Heatmap of the top 30 positively differentially expressed genes in clusters 2, 3, and 11 relative to all other clusters. Clusters were hierarchically clustered based on pseudobulk expression of the top 2000 variable genes. **j,** Dot plot showing expression of selected genes enriched in clusters 2, 3, and 11 compared to other clusters. Genes associated with chondrocyte differentiation are underlined in green, and genes enriched in tissue-specific chondrocyte expression are underlined in orange. Dot size reflects the percentage of cells expressing each gene; color intensity reflects mean normalized expression.

Grouping donors by high (BM D1 - BM D3) versus low (BM D4 - BM D6) osteogenic potential revealed minimal overlap in cell distributions (**Fig. 3b**). Single-cell trajectory analysis connected hBMSCs from donors with differing osteogenic potential, highlighting transcriptional distinctions along the inferred developmental path (**Fig. 3c**). The root of the inferred trajectory was anchored in BM D1-enriched cells, with low-performing donor cells occupying more distal pseudotime positions. This suggests that low-performing donors have diverged from the chondro-primed progenitor state rather than representing a more osteogenically committed population, consistent with their failure to develop bone *in vivo*. GO enrichment analysis of significantly upregulated genes in high osteogenic potential donors (adj. p-value < 0.05) revealed enrichment in ECM and cell-ECM interaction terms (**Fig. 3d**).

A more conservative pseudobulk analysis identified a restricted but biologically coherent set of genes enriched in high-potential donors (adj. p-value < 0.05, log2 fold change >1 or <–1, **Fig. 3e**). Rather than a broad transcriptional shift, these genes converge on the regulation of chondro-osteogenic commitment. Collagen type XI alpha 1 chain (COL11A1) encodes a fibrillar collagen essential for cartilage matrix integrity and endochondral bone development, with loss-of-function mutations causing severe skeletal dysplasias^48–50^. HAPLN1 stabilizes proteoglycan complexes in the pericellular matrix and promotes osteogenic differentiation through BMP/Smad signaling^51^. LIM and cysteine-rich domains-1 (LMCD1), a LIM domain transcription cofactor, similarly enhances osteogenic commitment downstream of BMP signaling^52^. Latent TGF-β binding protein (LTBP), by contrast, acts extracellularly to sequester latent TGF-β in the matrix, controlling its bioavailability and thus the balance between chondrogenic and osteogenic fate^53^. Altogether, that this gene set spans structural ECM components, intracellular signaling mediators, and extracellular growth factor regulators suggests these donors differ not in a single pathway but in the coordinated upstream state that primes the chondro-osteogenic axis.

Mapping these pseudobulk-enriched genes onto the UMAP showed their expression was not uniformly distributed but concentrated within three clusters (2, 3, and 11), suggesting these represent transcriptionally distinct subpopulations enriched for the chondro-osteogenic signature (**Fig. 3f and g**). Independent GO analysis confirmed that these clusters were specifically enriched for ECM-related cellular components relative to all remaining clusters (**Fig. 3h**), providing orthogonal support for their biological relevance. These clusters were therefore selected for deeper characterization. Differential expression analysis identified 78 genes significantly upregulated in these clusters relative to all others (log2 fold change > 2, adjusted p-value < 0.05), and pseudobulk profiling showed that they grouped together hierarchically, supporting their transcriptional relatedness (**Fig. 3i**). Chondrocyte differentiation emerged as a prominent enriched biological process. Consistent with this, TISSUES^54^ analysis identified chondrocytes as the top tissue association for the differentially expressed genes. A dot plot of significantly upregulated genes contributing to these GO annotations showed strong and selective enrichment within clusters 2, 3, and 11 (**Fig. 3j**). Together, these analyses identify chondrocyte-primed hBMSC subpopulations enriched in high potential donors, suggesting that donor-specific bone-forming capacity arises from intrinsic differences in cell state.

### TG-PEG hydrogels expose donor-specific differentiation capacity undetectable by conventional assays

We next asked whether the transcriptional divergence between donors is reflected in functional differences detectable by *in vitro* assays, and whether the matrix context required to reveal it *in vivo* is similarly necessary *in vitro*. First, to determine whether surface phenotype distinguishes high- from low-performing donors, we profiled surface marker expression as defined by the International Society for Cell & Gene Therapy (ISCT)^7^. All donors expressed mesenchymal-like markers and were negative for immune and endothelial markers (**Fig. 4a**). Despite some differences in CD49d and CD146 expression, hBMSCs from donors with high or low osteogenic potential were indistinguishable. To exclude the possibility that higher capacity to form bone *in vivo* was merely related to higher cell proliferation, we quantified hBMSC proliferation in 3D TG-PEG hydrogels for 16 days, which revealed equivalent proliferation across all donors (**Fig. 4b**). Thus, donor-specific differences in bone formation *in vivo* cannot be attributed to variations in canonical mesenchymal cell surface markers or proliferative capacity.

**Figure 4.**
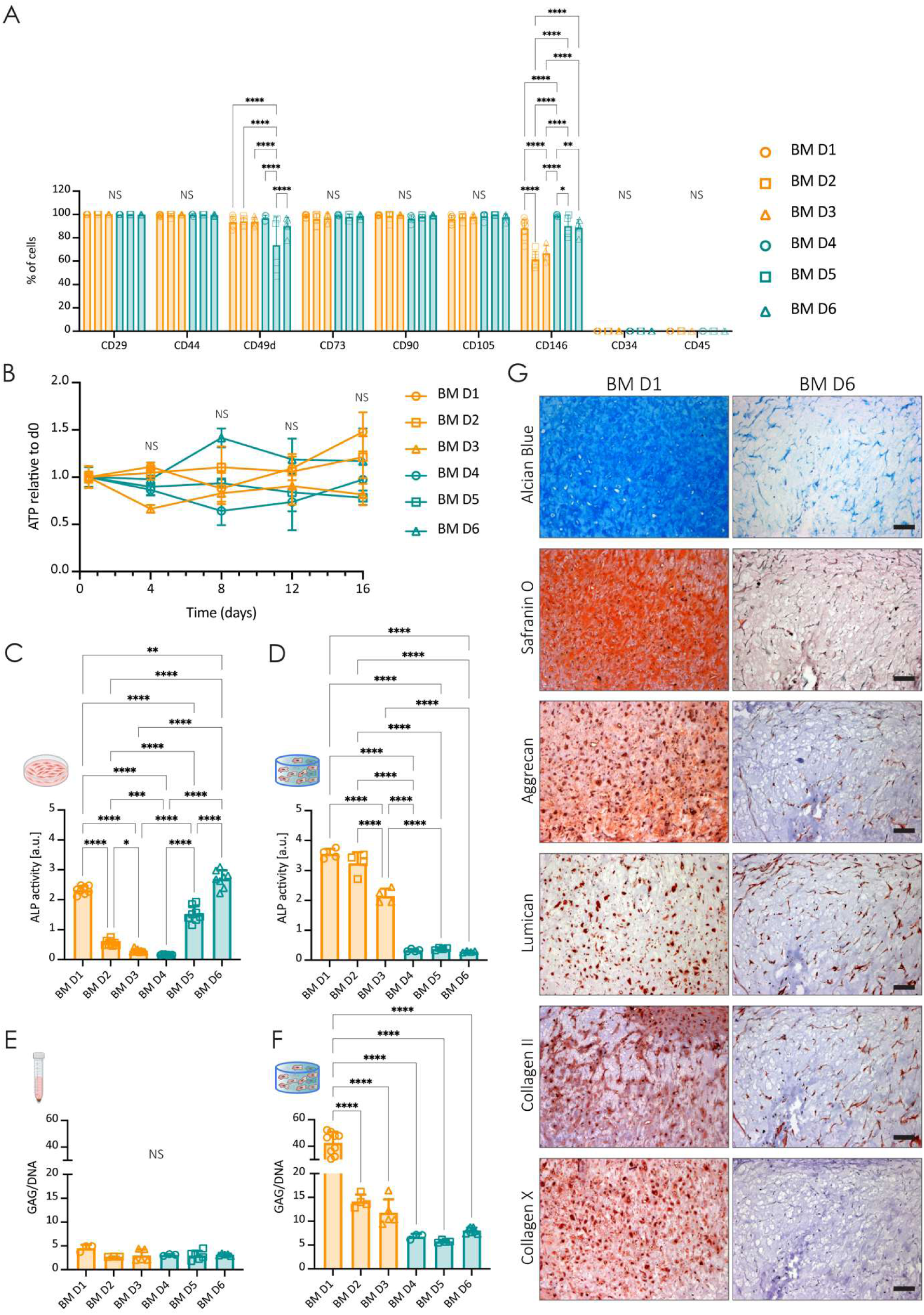
TG-PEG hydrogels expose donor-specific differentiation capacity undetectable by conventional *in vitro* assays. **a,** Flow cytometry analysis of consensus hBMSC surface markers. Expression of CD29, CD44, CD49d, CD73, CD90, CD105, and CD146, and absence of hematopoietic markers CD34 and CD45, were largely comparable across donors (N = 6 donors; n ≥ 4 independent experiments). **b,** Proliferation of hBMSCs from different donors cultured in 3D TG-PEG hydrogels (5 x 10^6^ cells per ml of hydrogel) up to 16 days *in vitro*, quantified by ATP content using the CellTiter-Glo assay (N = 6 donors; n = 5 hydrogels per time point and donor). Data are presented as mean ± s.d. For statistical analysis, to test whether proliferative capacity tracks with osteogenic performance, donors were grouped a priori by bone-forming potential as established in Fig. 2l (high-performing: BM D1-D3; low-performing: BM D4-D6). No significant difference in proliferation was detected between groups across the 16-day culture period (two-way ANOVA; group effect P = 0.5146, interaction P = 0.8640; Day 4: P > 0.9999, Day 8: P = 0.9986, Day 12: P = 0.9867, Day 16: P = 0.8043; Šídák’s multiple comparisons test). **c-g,** hBMSCs from different donors were cultured either in conventional cultures (monolayer or pellets) or encapsulated in 3D TG-PEG hydrogels and maintained under basal or lineage-specific differentiation conditions for 12 days for osteogenic or 21 days for chondrogenic differentiation. Schematics created with BioRender.com. **c,** Osteogenic differentiation potential of hBMSCs assessed in 2D monolayer, quantified by alkaline phosphatase (ALP) activity (N = 6 donors; n = 8 tissue culture wells per donor). **d,** Osteogenic differentiation potential of hBMSCs assessed in 3D TG-PEG hydrogels, quantified by ALP activity (N = 6 donors; n = 4 hydrogels per donor). **e,** Chondrogenic differentiation potential of hBMSCs evaluated in conventional pellet culture, quantified by glycosaminoglycan (GAG) deposition (N = 6 donors; n ≥ 3 pellets per donor). **f,** Chondrogenic differentiation potential of hBMSCs evaluated in 3D TG-PEG cultures, quantified by GAG deposition (N = 6 donors; n ≥ 3 hydrogels per donor). **g,** Representative histological and immunohistochemical staining of TG-PEG hydrogels containing hBMSCs from a high-performing (BM D1) and a low-performing (BM D6) donor following chondrogenic differentiation *in vitro*. Alcian blue and Safranin O staining visualize cartilage-rich matrix deposition; immunohistochemical staining for aggrecan, lumican, and collagen II marks chondrogenic differentiation, and collagen X marks hypertrophic maturation. Nuclei were counterstained with hematoxylin. Alcian blue stains sulfated glycosaminoglycans blue; Safranin O stains proteoglycan-rich matrix red. Scale bars: 100 µm. Bars represent mean ± s.d.; dots indicate independent experimental replicates (a) or intra-donor replicates (c-f). Two-way ANOVA with Tukey’s multiple comparisons test (a). Two-way ANOVA with Šídák’s multiple comparisons test (b). One-way ANOVA with Tukey’s multiple comparisons test (c-f). * P < 0.05; ** P < 0.01; *** P < 0.001; **** P < 0.0001; NS, not significant.

We therefore next asked whether intrinsic differentiation capacity might distinguish high- and low-performing donors. In conventional 2D osteogenic differentiation assays, alkaline phosphatase (ALP) activity did not correlate with *in vivo* bone-forming potential (**Fig. 4c** and controls in **Supplementary Fig. 3a**). Similarly, standard pellet-based chondrogenic cultures failed to discriminate between donors (**Fig. 4e** and controls in **Supplementary Fig. 3c**). In striking contrast, osteogenic differentiation of hBMSCs encapsulated in TG-PEG hydrogels partially recapitulated their *in vivo* performance (**Fig. 4d** and controls in **Supplementary Fig. 3b**). Chondrogenic stimulation within TG-PEG hydrogels resulted in significantly greater GAG deposition compared to pellet cultures (**Fig. 4f** and controls in **Supplementary Fig. 3d**), indicating enhanced chondrogenic maturation in hBMSCs within the 3D matrix. Unlike pellet cultures, TG-PEG conditions revealed clear functional differences between high- and low-osteogenic donors. These findings establish the TG-PEG microenvironment as a functional platform that exposes cell-autonomous donor-specific differentiation capacity undetectable in conventional culture systems and recapitulates *in vivo* bone-forming potential. Immunohistochemical analysis of 3D cultures confirmed enhanced chondrogenic differentiation in high (BM D1) versus low (BM D6) osteogenic potential donors. More specifically, Alcian blue and Safranin O stained sections showed a more pronounced cartilage matrix deposition by BM D1 than BM D6 (**Fig. 4g** and controls in **Supplementary Fig. 3e**). In line with these results, aggrecan, lumican, and collagen type II as well as, collagen type X, a hypertrophic marker, were significantly more deposited in hydrogels encapsulated with hBMSCs from BM D1 compared to BM D6 (**Fig. 4g** controls in **Supplementary Fig. 3f**). Collectively, these data establish that donor-specific osteogenic capacity *in vivo* is reflected in chondrogenic maturation *in vitro*, but only within a permissive 3D matrix context, raising the question of what intrinsic cellular program underlies this donor-to-donor divergence.

### Inherent ECM programs define high-performing hBMSC populations

This context-dependence suggested that reciprocal cell-matrix interactions, rather than intrinsic differentiation programming alone, govern donor-specific osteogenic performance. We therefore asked whether high- and low-performing donors differ in their capacity to establish distinct extracellular environments, independently of any differentiation stimulus. To test this directly, we exploited a key advantage of our fully synthetic platform. Unlike naturally derived matrices, the chemically defined and protein-free TG-PEG background enables unbiased proteomic interrogation of cell-deposited ECM without interference from exogenous matrix components. hBMSCs from BM D1 and BM D6 were therefore cultured in serum-free TG-PEG hydrogels for 14 days under basal conditions^26^, and the deposited matrisome was characterized by LC-MS/MS (**Fig. 5a** and **Supplementary Fig. 4a, b**). This experimental design interrogated the ECM program each donor intrinsically establishes, independent of any differentiation stimulus.

**Figure 5.**
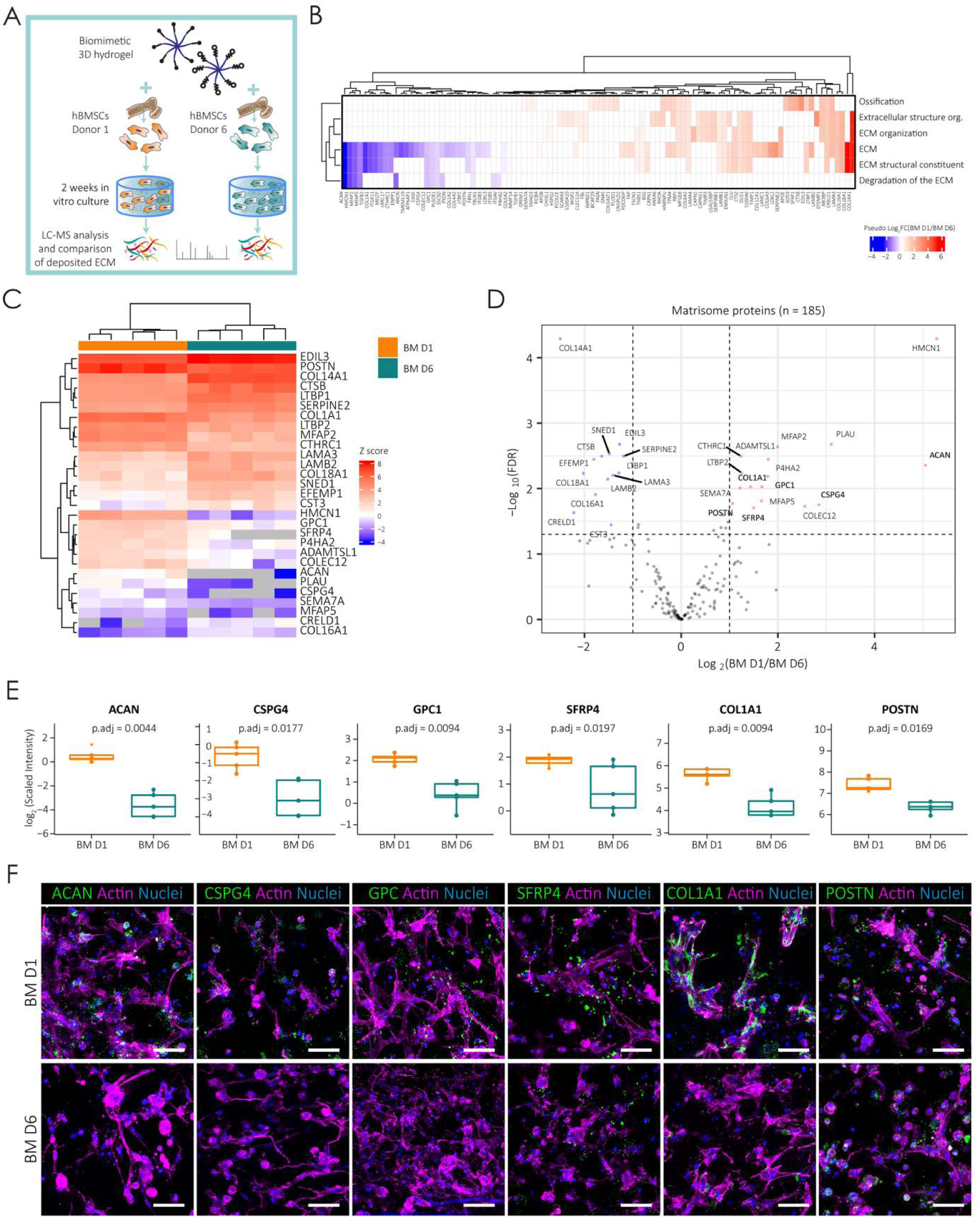
High-performing hBMSCs deposit a donor-specific pro-osteochondral ECM program revealed by unbiased proteomics. **a,** Schematic of the experimental workflow. hBMSCs (20 x 10⁶ cells ml⁻¹) from two donors (BM D1 and BM D6) were encapsulated in intermediate-stiffness TG-PEG hydrogels (500 Pa) and cultured *in vitro*. Cell-deposited extracellular matrix (ECM) was isolated and analyzed by liquid chromatography-mass spectrometry (LC-MS) after 14 days of culture (N = 2 donors; n = 5 hydrogels per donor). **b,** Enrichment maps highlight overrepresented pathways related to extracellular matrix organization and ossification in the selected modules (N = 2 donors; n = 5 hydrogels per donor). Protein co-expression modules were identified by weighted gene co-expression network analysis (WGCNA; **Supplementary** Fig. 4d). **c,** Unsupervised hierarchical clustering of the significantly up or down regulated ECM proteins, identified by cross-referencing with the Matrisome Project database^77,78^, reveals distinct ECM deposition profiles between donors BM D1 and BM D6 (N = 2 donors; n = 5 hydrogels per donor). **d,** Volcano plot showing differentially abundant ECM proteins deposited by hBMSCs from donors BM D1 and BM D6 following 14 days of *in vitro* culture in TG-PEG hydrogels. Proteins associated with cartilage and bone are highlighted in bold. The horizontal dashed line indicates the significance threshold (adjusted p < 0.05), and vertical dashed lines indicate fold-change thresholds (|log₂ fold change| ≥ 1) (N = 2 donors; n = 5 hydrogels per donor). **e,** Boxplots showing log_2_-scaled intensity of six ECM proteins identified as differentially enriched by proteomics (COL1A1, ACAN, GPC1, SFRP4, POSTN, and CSPG4) in TG-PEG hydrogels containing hBMSCs from BM D1 and BM D6 after 14 days of culture under basal conditions (N = 2 donors; n = 5 hydrogels per donor). Unpaired two-tailed Student’s t-test. **f,** Validation of proteomic findings by immunofluorescence staining of whole TG-PEG hydrogels cultured *in vitro*. Representative images show ECM proteins (green) including aggrecan (ACAN), chondroitin sulfate proteoglycan 4 (CSPG4), glypican-1 (GPC), secreted frizzled-related protein 4 (SFRP4), collagen type I alpha 1 chain (COL1A1), and periostin (POSTN); the actin cytoskeleton (magenta), and nuclei (DAPI, blue) for donors BM D1 and BM D6. Scale bars: 50 µm.

Unsupervised clustering analysis showed donor-based clustering of samples (**Supplementary Fig. 4c**). Weighted gene co-expression network analysis (WGCNA) identified distinct protein modules that separated BM D1 and BM D6 (**Supplementary Fig. 4d**). Enrichment maps for overrepresented pathways related to ECM and ossification in the selected modules (**Fig. 5b**) revealed a clear segregation of BM D1 and BM D6 based solely on their deposited ECM composition, indicating that these donors establish fundamentally distinct extracellular environments (**Fig. 5c**).

Inspection of the most differentially enriched matrisome proteins in BM D1-derived matrices revealed a coherent chondro-osteogenic progression signature rather than a collection of unrelated hits (**Fig. 5d, e**). Aggrecan (ACAN) and chondroitin sulfate proteoglycan 4 (CSPG4) are core structural components of cartilage matrix that provide the glycosaminoglycan-rich scaffold required for chondrocyte differentiation and early endochondral template formation^55^. Glypican-1 (GPC1), a heparan sulfate proteoglycan, modulates local growth factor gradients and has been linked to osteogenic niche patterning^56^. Secreted frizzled-related protein 4 (SFRP4) acts as an extracellular modulator of Wnt signaling, with established roles in regulating the chondrocyte-to-osteoblast transition during endochondral ossification^57^. Collagen type I alpha 1 chain (COL1A1) and periostin (POSTN) mark the later stages of this program: COL1A1 as the primary structural collagen of mineralizing bone matrix^58^, and periostin as a periosteum-associated matricellular protein that supports osteoblast differentiation and cortical bone formation^59^. This signature underlying endochondral progression emerges under basal conditions within the TG-PEG system, in the absence of exogenous biochemical inductive cues, suggesting it reflects an intrinsic property of high-performing donors rather than a response to exogenous stimulation.

Immunofluorescence analysis of whole TG-PEG hydrogels validated the increased deposition of all six proteins in BM D1 relative to BM D6 (**Fig. 5f**), confirming the proteomic signature. Critically, this is not merely a transcriptional difference between donors, high-performing hBMSCs actively translate their chondro-primed cell state into a compositionally distinct extracellular environment at the protein level, establishing an ECM niche consistent with the molecular architecture of the endochondral template. Moreover, because this ECM program emerges under basal conditions *in vitro*, in the absence of any exogenous differentiation stimulus or *in vivo* context, it reflects an intrinsic property of high-performing donors rather than an adaptive response, suggesting that osteogenic competence is encoded at the level of cell state before any commitment signal is received. That cell-deposited ECM is itself a functional determinant of downstream tissue formation is consistent with emerging work showing that hBMSC-derived matrix carries instructive properties for musculoskeletal regeneration^60^. Together, these data identify intrinsic ECM remodeling capacity as a defining correlate of donor-specific osteogenic performance within the TG-PEG context, and raise the question of whether this *in vitro* molecular program translates into a distinct developmental trajectory *in vivo*.

### Longitudinal *in vivo* tracking reveals a donor-specific endochondral cascade driving bone formation

The proteomics data established that high- and low-performing donors deposit fundamentally distinct ECM programs *in vitro*. To determine whether this molecular divergence translates into different developmental fates *in vivo*, we first confirmed that the proteomic ECM signature identified *in vitro* is enacted *in vivo*. Indeed, BM D1 ossicles showed stronger expression of all identified matrisome markers compared to BM D6 at 2 weeks post-implantation (**Supplementary Fig. 4e** and **Fig. 6b**). We then tracked BM D1 and BM D6 constructs longitudinally at 1, 2, and 4 weeks post-implantation to resolve the sequential events distinguishing productive endochondral ossification from fibrous tissue deposition.

**Figure 6.**
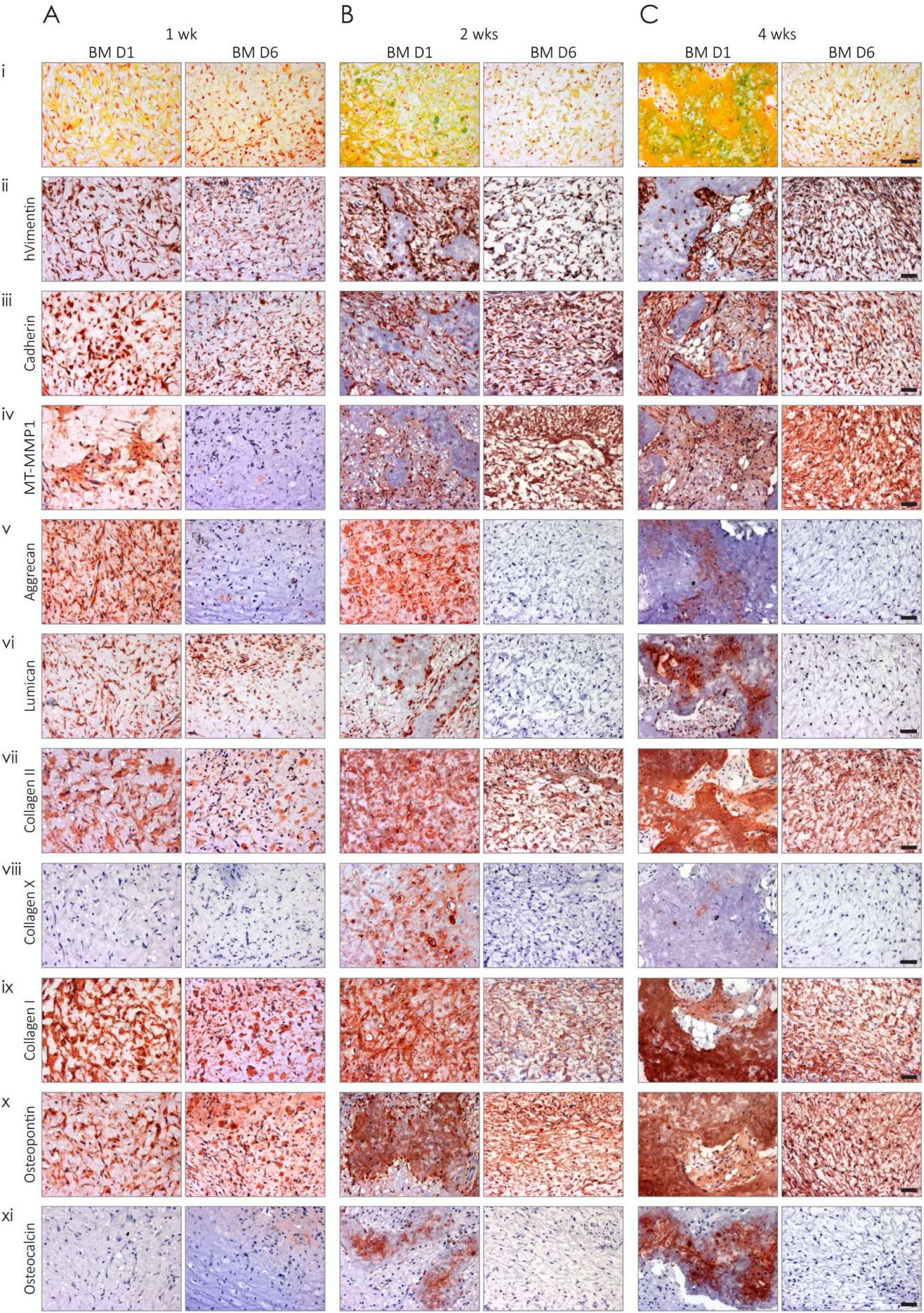
Longitudinal *in vivo* tracking reveals a donor-specific endochondral ossification cascade driving bone formation. hBMSCs from a high-performing donor (BM D1) or a low-performing donor (BM D6) were encapsulated in optimized intermediate-stiffness TG-PEG hydrogels containing RGD adhesion motifs and MMP-sensitive degradable sites and implanted subcutaneously in immunodeficient mice. **a-c,** Representative histological and immunohistochemical analysis of implanted constructs at 1 week (**a**), 2 weeks (**b**), and 4 weeks (**c**) post-transplantation. **i)** Modified MP staining illustrates temporal changes in extracellular matrix composition, including glycosaminoglycan-rich cartilage matrix, collagenous bone matrix, and vascularized tissue. Immunostaining highlights progressive stages of tissue development, including **iii)** cell-cell condensation (cadherin), **iv)** matrix remodeling (MT-MMP1), **v-vii)** chondrogenic differentiation (aggrecan, lumican, and collagen II), **viii)** hypertrophic maturation (collagen X), and **ix-xi)** osteogenic differentiation (collagen I, osteopontin, and osteocalcin). Human cells were identified using **ii)** human-specific vimentin staining. Nuclei were counterstained with hematoxylin. Across time points, constructs containing BM D1 hBMSCs display early extracellular matrix deposition, acquisition of chondrocyte-like morphology, and subsequent progression toward mineralized bone tissue, whereas constructs containing BM D6 hBMSCs show limited matrix deposition and lack of coordinated endochondral progression. Scale bars: 50 µm (all panels). In MP staining, glycosaminoglycan-rich cartilage appears blue/green, collagenous bone matrix appears yellow, vascularized tissue and marrow appear red, and nuclei appear black.

MP staining revealed progressive morphological and matrix changes consistent with endochondral ossification in BM D1 constructs (**Fig. 6a-c**). As early as one week post-implantation, hydrogels containing BM D1 hBMSCs showed GAG deposition, which intensified at weeks 2 and 4 (**Fig. 6i**). By week 2, embedded cells adopted a rounded, chondrocyte-like morphology and organized into condensations that subsequently transitioned into mineralized bone structures by week 4. In contrast, constructs with BM D6 hBMSCs lacked substantial GAG deposition and failed to undergo morphological progression toward a chondrogenic intermediate. Human-specific vimentin staining confirmed persistence of both BM D1 and BM D6 hBMSCs within implants (**Fig. 6ii**), excluding differential cell survival as a contributing factor.

Consistent with the *in vitro* differentiation and proteomic signatures, BM D1 constructs within TG-PEG hydrogels displayed sequential activation of developmental markers characteristic of endochondral ossification, including cadherin-mediated cell condensation (**Fig. 6iii**), matrix remodeling via membrane-type matrix metalloproteinase 1 (MT-MMP1) (**Fig. 6iv**), chondrogenic markers (**Fig. 6v-vii**), hypertrophic differentiation (**Fig. 6viii**), and subsequent osteogenic maturation (**Fig. 6ix-xi**). Together, these findings demonstrate that the defined TG-PEG microenvironment enables high-performing hBMSCs to spontaneously undergo a coordinated endochondral ossification program *in vivo*, recapitulating hallmark features of embryonic endochondral bone development^61^, whereas low-performing donors fail to initiate this developmental cascade.

## Outlook

Collectively, these results demonstrate that human skeletal progenitor competence resides within intrinsic cell states that are only revealed within a defined synthetic matrix context, and most completely within a permissive, exogenous growth factor-free niche that permits cell-intrinsic programs to operate without interference. By decoupling extrinsic osteoinductive signals from cell-intrinsic behavior, the TG-PEG platform uncovers that donor-specific bone-forming capacity is functionally enacted through a coordinated pro-osteochondral ECM deposition program that drives spontaneous endochondral ossification *in vivo*. This program is rooted in chondrocyte-primed transcriptional states present before any commitment signal is received. Remarkably, scRNA-seq and proteomic data converge independently on the same chondro-osteogenic axis at transcriptional and protein levels, respectively, suggesting a shared underlying program. Critically, BMP-2 supplementation actively overrides this intrinsic program, collapsing inter-donor variability in bone-forming capacity and masking functional differences. Thus, the TG-PEG microenvironment functions not merely as a scaffold but as a functional interrogation tool through which intrinsic competence becomes legible. These findings have potential implications for how the field evaluates and selects skeletal progenitors for therapeutic use^18,19,42,43,62^.

The chondrocyte-primed hBMSC subpopulations identified here provide a strong transcriptional framework for future studies. Whether prospective isolation of these subpopulations (using surface markers derived from their transcriptional signatures) yields cells with uniformly high osteogenic performance remains to be established. It also remains to be determined whether these subpopulations are present and similarly predictive in freshly isolated, non-passaged hBMSCs, with direct implications for clinical translation. Additionally, while six donors provided sufficient power to detect robust functional differences, expansion to larger and more clinically diverse cohorts will be required to establish the generalizability of ECM-based donor stratification. It is important to note that the functional stratification of donors demonstrated here was established exclusively within the TG-PEG platform. Whether the distinction between high- and low-performing donors reflects an absolute difference in osteogenic potential (or a differential capacity to interface with this specific synthetic matrix) cannot be fully resolved without validation in independent delivery systems. The convergence of transcriptional differences identified by scRNA-seq in 2D-expanded cells with the functional and proteomic differences observed in TG-PEG hydrogels provides indirect evidence for cell-intrinsic rather than matrix-specific divergence, but direct validation using orthogonal biomaterial platforms or in vivo transplantation assays without matrix will be required to confirm this interpretation. Similarly, the ECM components identified by proteomics represent correlates of high osteogenic performance within this context; their causal roles remain to be established through perturbation experiments.

The chondrocyte-primed ECM program identified here also raises a question with immediate therapeutic relevance: can low-performing donors be rescued? Our data suggest that the divergence between high- and low-performing donors is cell intrinsic rather than imposed by the TG-PEG environment specifically, but this does not preclude the possibility that matrix mechanical, biophysical or biochemical cues^63,64^ could promote entry into the chondro-primed state in low-potential cells. The modular tunability of TG-PEG hydrogels makes them well suited to screen such interventions systematically.

Our data further suggest that the intrinsic cell state predicting bone formation also determines hematopoietic niche establishment. Only high-performing donors generated substantial bone volume alongside a functional hematopoietic niche, and these were the same donors that most clearly progressed through the endochondral program. This link between endochondral competence and marrow niche establishment aligns with prior work demonstrating that endochondral ossification is required for hematopoietic stem cell niche formation^46^, and suggests that donor stratification based on intrinsic ECM programs may be relevant not only for bone repair but for engineering functional human ossicles as models of hematopoietic disease and marrow biology.

More broadly, this work establishes that synthetic matrices designed for permissiveness rather than instruction can serve as resolution tools to uncover functional heterogeneity in human stem and progenitor cell populations otherwise masked by conventional culture conditions. While demonstrated here for skeletal progenitors, this approach may be applicable to other stem cell systems where intrinsic competence is similarly obscured by exogenous instructive cues, offering a generalizable strategy to functionally interrogate human cell potential in the absence of biological noise.

## Methods

All chemicals and materials were purchased from Sigma Aldrich or Thermo Fisher Scientific unless otherwise specified.

### Animal care

All animal procedures were approved by the veterinary offices of the Swiss cantons Zürich and Lausanne under the ethical license (Application No. ZH169/2015). Experiments and handling of mice were conducted in accordance with the Swiss law of animal protection. 6-8-week-old immunodeficient HsdCpb:NMRI*-Foxn1^nu^* (nude) female mice (purchased from Envigo) or NOD.Cg-*Prkdc^scid^Il2rg^tm1Wjl^/*SzJ (NSG) female mice (purchased from The Jackson Laboratory) were used for the following experiments.

### Human cell isolation and culture expansion

Human bone marrow-derived stromal cells (hBMSCs) were isolated as previously described^65^ from bone marrow aspirates of healthy donors (n = 6) obtained during orthopedic surgical procedures after informed consent and in accordance with the local ethical committee (University Hospital Basel; Prof. Dr. Kummer; approval date 26/03/2007 Ref Number 78/07). Age of healthy donors ranged from 17 to 39. Cells were cultured at 37 °C in a humidified atmosphere at 5% CO_2_ in minimum essential medium alpha (MEMα, with nucleosides, Gibco) supplemented with fetal bovine serum (FBS, 10%, Gibco), penicillin (100 U ml^-1^, Gibco), streptomycin (100 µg ml^-1^, Gibco) also referred as 1% P/S, and fibroblast growth factor-2 (FGF-2, 5 ng ml^-1^, PeproTech). Cells were passaged before reaching 90% confluency, and medium was changed every 2-3 days. For hydrogel encapsulation, cells were trypsinized and resuspended in 10% FBS/MEMα in the desired concentration and mixed in the corresponding precursor mixes.

### TG-PEG precursors synthesis and preparation

TG-PEG precursors were synthesized as previously described^23,66^. Briefly, 8-arm PEG-VS (PEG-vinylsulfone, 40 kDa MW; NOF) was functionalized with peptides (obtained with a purity of > 95% from Bachem AG) that contained an earlier described cysteine cassette (ERCG) optimized for its reaction with PEG-VS and either a factor XIII (FXIII) glutamine acceptor substrate sequence (Gln; H-NQEQVSPL-ERCG-NH_2_) or a matrix metalloproteinase-degradable (*in italics*) lysine donor substrate (MMP_sensitive_-Lys; Ac-FKGG-*GPQGIWGQ*-ERCG-NH_2_), or instead a non-degradable lysine donor substrate (ND-Lys; Ac-FKGG-GDQGIAGF-ERCG-NH_2_) for non-degradable TG-PEG gels. A 1.2 molar excess of peptides over PEG-VS was reacted in triethanolamine (TEA) at pH 8.0 for 2 h at 37 °C. Resulting 8-PEG-Gln and 8-PEG-MMP_sensitive_-Lys or 8-PEG-ND-Lys precursors were excessively dialyzed against pure water, lyophilized and stored at -20 °C until further use. The crosslinking enzyme, plasma-derived transglutaminase FXIII (200 U ml^−1^, Fibrogammin P, CSL Behring), was activated with thrombin (2 U ml^−1^) for 30 min at 37 °C and stored in small aliquots at -80 °C until use.

### 3D TG-PEG hydrogel formation

Stoichiometrically balanced solutions of 8-arm PEG-Gln and 8-arm PEG-MMP_sensitive_-Lys, or 8-arm PEG-Gln and 8-arm PEG-ND-Lys for non-degradable gels were prepared in Tris buffer (50 mM, pH 7.6) containing calcium chloride (CaCl_2_, 50 mM). Additionally when stated, 50 µM Lys-RGD peptide (Ac-FKGG-RGDSPG-NH_2_), indicated amounts of cells, 12.5 ng µl^-1^ BMP-2 (produced as previously described^67^), or indicated amounts of hydroxyapatite tricalcium phosphate (HA/TCP) particles (60/40, diameter < 75 µm, CAM Bioceramics) were added to the precursor solution. To immobilize recombinant human Jagged1–Fc protein (R&D Systems, 1277-JG-050) or recombinant human Delta-like-4 (DLL4)–Fc protein (Sino Biological, 10171-H02H) into TG-PEG hydrogels, these ligands (500 nM final concentration) were pre-incubated for 30 min with Gln-ZZ protein (5 µM final concentration, previously characterized^41^) to pre-form the Gln-ZZ/Fc-protein complex, and added to the gel precursor mix. Subsequently, hydrogel crosslinking of final dry mass content of 0.9% (corresponding to low stiffness), 1.7% (for intermediate stiffness) or 5% (for high stiffness) (w/v) was initiated by the addition of 10 U ml^-1^ of activated transglutaminase factor XIII, followed by vigorous mixing. Disc-shaped matrices were prepared between hydrophobic glass slides (treated with SigmaCote) and incubated for 30 min at 37 °C in a humidified atmosphere at 5% CO_2_. After completed polymerization, hydrogels were released from glass slides and transferred to tissue-culture plates for *in vitro* experiments or stored in a humidified atmosphere for immediate *in vivo* implantation.

### Natural matrices

hBMSCs (at final concentration of 20x10^6^ cells per ml of hydrogel) were mixed in cold solutions of 3 mg ml^-1^ of collagen (PureCol EZ Gel solution, Sigma) or ECM-like matrix (referred as ECMatrix in the text, ECM625, Merck Millipore) following manufacturer’s instructions. For fibrin gels as previously described^68^, hBMSCs (20x10^6^ cells per ml of hydrogel) were added to a solution of 4 mg ml^-1^ of fibrinogen containing 2.5 mM CaCl_2_. Next, FXIII (2 U ml^-1^) and thrombin (2 U ml^-1^) were incorporated to begin hydrogel crosslinking. Cell-laden hydrogels were then incubated for 30 min at 37 °C to enable complete polymerization prior to immediate (within 2 hours) *in vivo* implantation.

### Hydrogel stiffness characterization by rheometry

Hydrogel gelation was analyzed on a rheometer (MCR 301, Anton Paar) equipped with 20 mm plate–plate geometry (PP20, Anton Paar) at 37 °C in a humidified atmosphere. For *in situ* measurements gel mixtures were precisely loaded onto the center of the bottom plate. The upper plate was lowered to a measuring gap size of 0.2 mm, ensuring proper loading of the space between the plates and gel precursors, the dynamic oscillating measurement was then started. The evolution of storage modulus (G’) and loss modulus (G’’) at a constant angular frequency of 1 Hz and constant shear strain of 4% was recorded for 30 min when equilibrium was reached. For measurements of TG-PEG hydrogels containing HA/TCP particles, to avoid particle precipitation during the rheological measurements, hydrogels were pre-formed. Hydrogels were prepared 24 h before measurements and incubated in Tris buffer at 37 °C. Swollen hydrogels were then loaded onto the rheometer, compressed by 10% and measured with a frequency sweep at 1% strain.

### Subcutaneous implantation of humanized hydrogels in mice

To create subcutaneous pouches in nude mice ca. 6 mm lateral skin incisions were made in four dorsal positions. Following randomization, pre-formed 18 µl gelled 3D scaffolds containing hBMSCs (20x10^6^ per ml of hydrogel, unless indicated) were placed in these pouches. Incisions were closed with 6.0 Vicryl sutures (Ethicon) and animals were closely monitored. One, two, four or eight weeks post-implantation unless stated otherwise, animals were euthanized via CO_2_ asphyxiation, ossicles and specified murine organs were retrieved and immediately fixed in 4% (v/v) formalin solution for 12 h at 4 °C. After fixation they were washed in PBS and analyzed by microCT and histologies.

### Micro-computed tomography analysis at endpoints

After fixation in formalin and storage in PBS, ossicles were scanned in a microCT 40 (Scanco Medical AG) operated at an energy of 55 kVp and intensity of 72 µA at 4 W. Scans were executed at a high-resolution mode resulting in a voxel size of 10 µm. In reconstructed images bone tissue was segmented from background using a global threshold of 10% of maximum grey value. A cylindrical mask with a diameter of 5 mm was manually placed around the ossicle. Bone volume within the mask was measured using the ImageJ plugin BoneJ^69^. Reconstructions were assembled using the 3D Viewer plugin in ImageJ.

### Tissue processing and histological staining

Next, samples were decalcified for 2 to 6 weeks in 10% EDTA solution with continuous shaking at 4 °C, followed by paraffin embedment and microtome sectioning at 4 µm. Sections were deparaffinized and rehydrated through graded ethanol solutions prior to hematoxylin and eosin (H&E), Alcian blue, Safranin O, and modified Movat’s pentachrome (MP)^70^ staining using standard histological procedures. Alcian blue and Safranin O staining were used to assess glycosaminoglycan- and cartilage-rich extracellular matrix deposition, respectively. Briefly for MP staining, sections were sequentially incubated with alcian blue (Fluka, 1 g in 100 ml 1% glacial acetic acid) for 10 min, alkaline alcohol (10% ammonium hydroxide in 95% EtOH) for 1 hour, Weigert’s iron hematoxylin (Weigert reagent A and B in a ratio 1:1) for 20 minutes, brilliant crocein R-fuchsine (1 part Biebrich Scarlet/Acid Fuchsin solution and 1 part 0.2% acidic fuchsin in 0.5% acetic acid) for 20 minutes, 0.5% acetic acid for 30 seconds, 5% phosphotungstic acid (PWS) for 15 min, 0.5% acetic acid for 2 minutes, washed 3 times with absolute ethanol for 5 minutes, 10 min in safran dye (6 g safran powder in 100 ml absolute EtOH), dehydrated and mounted in mounting medium. In MP, red (fibrin) indicates muscle/vascularized tissue; yellow (reticular fibers/collagen) indicates bone; green/blue (mucin) indicates cartilaginous tissue; and black, nuclei and elastic fibers. Stained sections were dehydrated, mounted, and imaged using a Zeiss 200M inverted microscope.

### Immunohistochemistry (IHC) and immunofluorescence (IF) stainings

For IHC or IF, slides were deparaffinized and rehydrated. Next, different antigen retrievals were performed based on the needs of each staining. Collagen stainings required enzymatic digestion with hyaluronidase (2 mg ml^-1^ in PBS) for 45 min, followed by 30 min incubation in pronase (1 mg ml^-1^ in PBS). For the rest of stainings heat antigen retrieval was performed, sections were incubated for 20 min at 90 °C in 10 mM sodium citrate, 0.05% Tween-20 at pH 6.0 or in 10 mM Tris Base, 1 mM EDTA at pH 9.0. All sections were then blocked for 1 h at room temperature in 1.5% bovine serum albumin (BSA) 0.5% Tween-20 in PBS, followed by primary antibody incubation overnight at 4 °C in PBS containing 1% BSA in a humidified chamber. Next day, sections were incubated with the corresponding secondary antibody in PBS containing 1% BSA for 1 h at room temperature in the dark. Between individual incubation steps, sections were rinsed three times with PBS containing 0.1% Tween-20. IF slides were washed and mounted with fluorescent mounting medium containing 40,6-diamidino-2-phenylindole (DAPI, Abcam, ab104139). IHC slides were developed using the peroxidase substrate ImmPACT AMEC Red (Vector Laboratories, SK-4285), counterstained with hematoxylin (Vector Laboratories, H-3404), and mounted in aqueous mounting media. **Table 1** details the employed antibodies and their concentrations. For IF stainings, inverted laser-scanning microscope Leica TCS SP5 was used. For IHC, images were acquired using a Zeiss 200M inverted microscope. Images were further processed in Fiji. Negative controls, consisting of either omission of the primary antibody or isotype-matched controls, were included in all staining experiments.

**Table 1.**
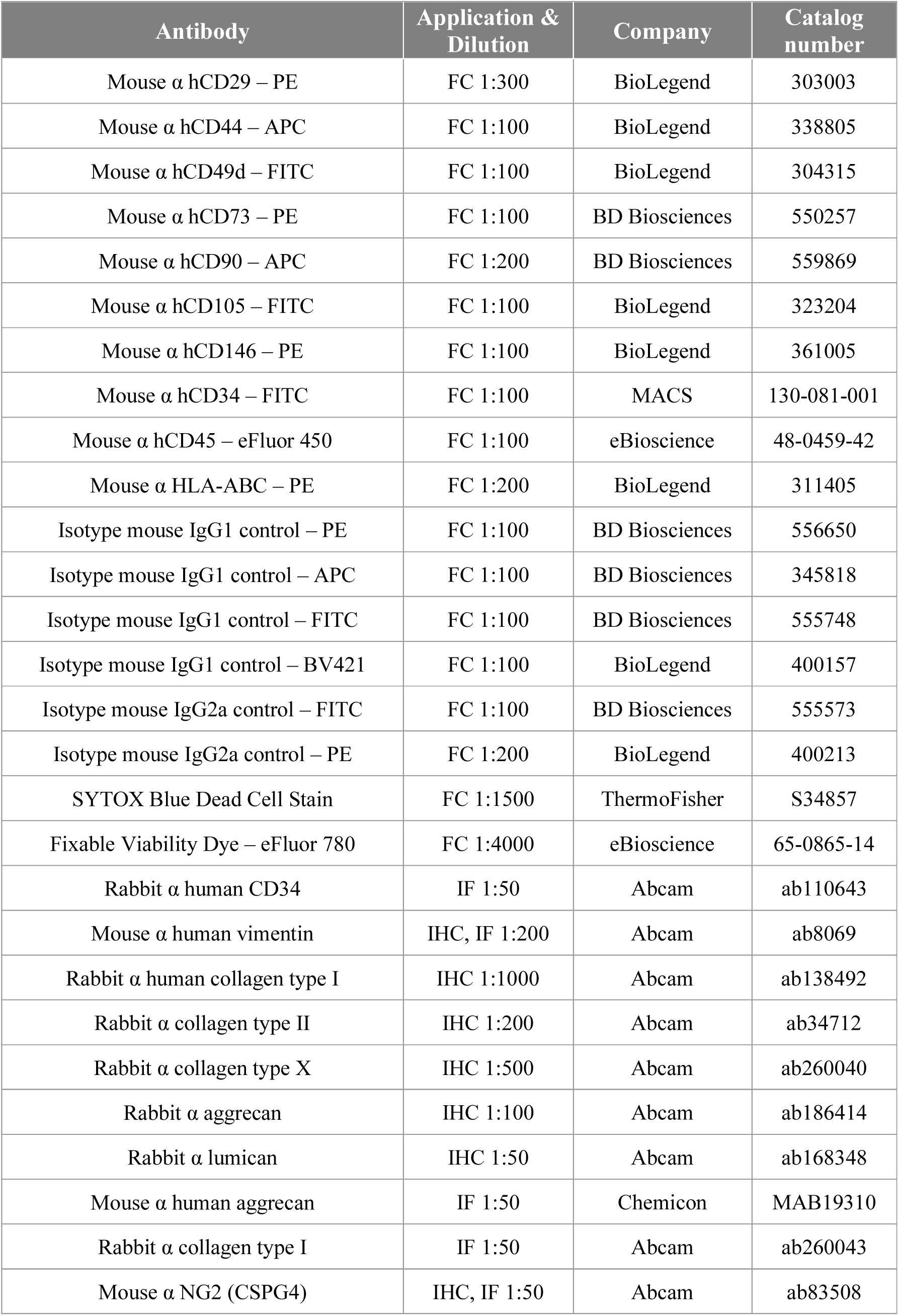
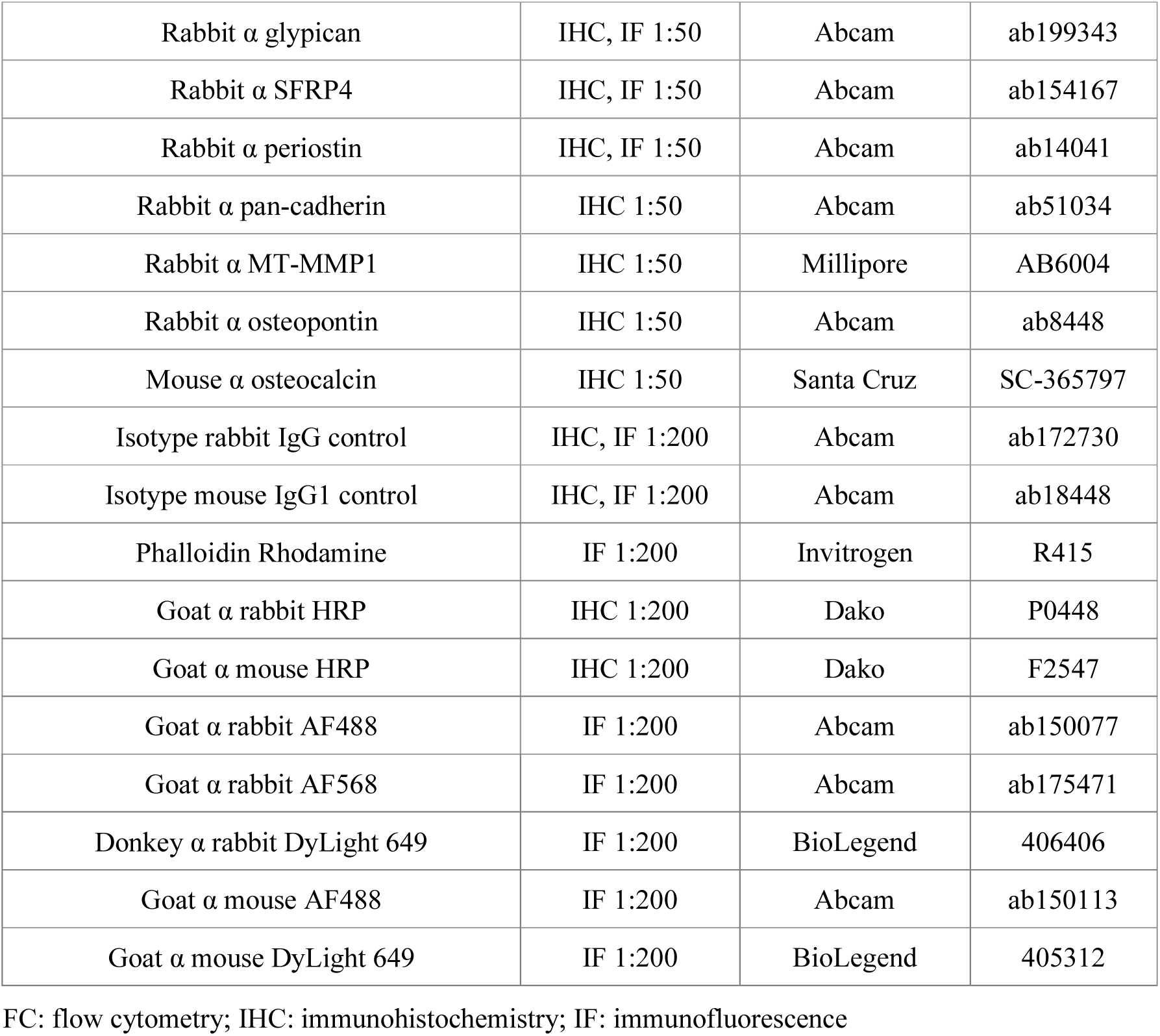
List of detailed antibodies used for flow cytometry and histological stainings.

### NSG hematopoietic xenotransplantation

Primary human CD34^+^ cells were isolated from umbilical cord blood collected from healthy donors at the University Hospital Zürich after obtaining informed consent. The study was approved by the ethics board of the canton Zürich, Switzerland (approval date 21/03/2007; Ref Number 07/07). Blood (typically between 30-70 ml) was subsequently centrifuged to enrich for mononuclear cells, which were in turn further magnetically sorted for CD34^+^ cells using positive immunomagnetic selection (CD34^+^ MicroBead Kit, Miltenyi Biotec) according to the manufacturer’s instructions obtaining > 80% CD34^+^ cells. Cells were directly frozen down until use. Upon NSG transplantation, cells were thawed in 50% IMDM: 50% FBS, centrifuged 5 min at 300 g and subsequently resuspended in PBS. Then, NSG mice that had 8 weeks before received a subcutaneous implantation of hBMSC-laden TG-PEG scaffolds, received 0.6x10^6^ human CD34^+^ cells via tail vein injection (200 µl per animal). Animals were sacrificed 16 weeks post-CD34^+^ human cells transplantation. Ossicles and organs were excised and immediately fixed in 4% (v/v) formalin solution for 12 h at 4 °C. After fixation they were washed in PBS and further analyzed by microCT and histologies.

### Calvarial defect healing model

Optimized TG-PEG hydrogels of intermediate stiffness (1.7% w/v, containing MMP-degradable and RGD sites) were loaded with hBMSCs (20x10^6^ per ml of hydrogel), crosslinked into 11 µl discs sized to fill the calvarial defect, and subsequently implanted within 2 hours of gelation. Craniotomies of 4 mm diameter were created in the parietal bones of the skull, one on each side of the sagittal suture. Pre-formed hydrogel discs were placed in the cranial defects and the skin was closed with 6.0 Vicryl sutures (Ethicon). Mice were euthanized 4 weeks post-implantation, calvaria were excised and immediately fixed in 4% (v/v) formalin solution for 12 h at 4 °C. After fixation they were washed in PBS and further analyzed by microCT and histologies.

### Osteogenic pre-differentiation of hBMSCs prior to implantation

To assess the effect that osteogenic differentiation had on hBMSC capacity to form bone *in vivo*, hBMSCs were cultured in 2D or in 3D optimized TG-PEG hydrogels (18 µl hydrogels, 1.7% TG-PEG containing MMP-degradable and RGD sites, with 20x10^6^ cells per ml of hydrogel) with osteogenic medium composed of MEMα (with nucleosides), FBS (10%), penicillin (100 U ml^-1^), streptomycin (100 µg ml^-1^), 10 mM HEPES, 1 mM sodium pyruvate, 2 mM L-glutamine, 50 µg ml^-1^ L-ascorbic acid, 10 mM β-glycerol phosphate, and 100 ng ml^-1^ BMP-2. Medium was replaced every 3 days. After 8 days, pre-differentiated hBMSCs in 2D were encapsulated in optimized TG-PEG hydrogels (18 µl, 20x10^6^ cells per ml of hydrogel) and immediately (within 2 hours) subcutaneously implanted in nude mice. For 3D pre-differentiated hBMSCs, gels were washed in PBS, and subsequently implanted. hBMSCs that had not been pre-differentiated were also encapsulated and implanted as previously described to serve as positive controls.

### Scanning electron microscopy

For scanning electron microscopy (SEM), TG-PEG hydrogels containing HA/TCP particles were fixed in 50% osmium in phosphate buffered saline (PBS). After PBS washing, they were dehydrated by 30-minute period in 70% ethanol, followed by 80% and 100% ethanol and subsequently treated with hexamethyldisilazane (HMDS) for 1 hour, and then dried on air overnight. Lastly, samples were platinum sputter coated. Imaging was performed on a Zeiss Supra 50 VP at the UZH Center for Microscopy and Image Analysis (ZMB).

### Isolation and flow cytometry sorting of implanted ossicles

Scaffolds were excised from the subcutaneous pockets at one week post-surgery and collected in MEMα + 10% FBS on ice. Connective tissue around the ossicles was carefully removed. Next, ossicles were first mechanically dissociated into small pieces in Eppendorf tubes, followed by addition of digestion medium (MEMα + 10% FBS + 1 mg ml^-1^ Collagenase A (Roche 11088793001) + 0.5 mg ml^-1^ DNase I (Roche 11284932001)), and were incubated at 37 °C rocking for 1 h. Followed by trypsin (final conc. 0.025 %, Gibco) addition for 10 min at 37 °C, cells were washed with FACS buffer (PBS + 1 mM EDTA + 2% FBS) and filtered through a 70-µm cell strainer (BD Falcon 352350). Red blood cells were lysed by incubation with RBC lysis buffer (Biolegend 420301) for 5 min at room temperature and cells were washed and re-suspended in FACS buffer. Next, cells were incubated for 10 min at 4 °C with mouse Fc block solution (1μg per 10^6^ cells in 100 μl, BD Biosciences 553141) prior to receiving the antibody mix solution. To sort all human cells, despite their surface markers or differentiation state, a general anti-human HLA-ABC antibody (**Table 1**) was used. Cells were stained in FACS buffer for 30 min rocking at 4 °C in the dark. Cells were then washed and resuspended in FACS buffer to be analyzed and sorted in BD FACSAria III. Dead cells were stained by the Fixable Viability Dye and were, alongside doublets, excluded from analysis. Gates were defined according to the fluorescence intensity of the isotype control and fluorescence minus one (FMO) staining. Next, human cells were sorted through a 100 µm nozzle with 20 psi sheath pressure directly into RNA lysis buffer for further bulk RNA-sequencing analysis.

### RNA isolation and bulk mRNA sequencing

RNA was isolated from sorted hBMSCs retrieved from *in vivo* ossicles one week post-implantation or retrieved from hydrogels cultured *in vitro* (named hBMSCs non-implanted to serve as controls) following the Direct-zol RNA MiniPrep (Zymo Research, R2052). RNA sequencing libraries were prepared as previously described using BRB-seq method^44^. Data analysis and gene ontology enrichment was performed using the ASAP platform^71^. Briefly, genes were filtered based on count per million (CPM, 1 CPM in minimally 1 sample detected), followed by Voom normalization^72^. Differential expression was done using limma with adjusted p-value (FDR) ≤ 0.05 and fold change (FC) ≥ 2.

### Single cell RNA-sequencing

hBMSCs were expanded in vitro and detached using trypsin prior to fixation with the Parse Biosciences Fixation Kit (v3). Sample barcoding, complementary DNA (cDNA) amplification, and library preparation were performed following the Parse Biosciences Evercode WT V2 protocol. Libraries were sequenced on an Illumina NovaSeq X Plus platform, and resulting FASTQ files were processed using the Trailmaker™ pipeline. Quality control was performed in Trailmaker, with minimum transcript thresholds applied automatically. Mitochondrial content and doublet filters were also set to automatic thresholds. Filtered results were exported as a Seurat object.

Data normalization and dimensionality reduction were conducted using Seurat (v5.3.0). Counts were normalized using the LogNormalize method with a scale factor of 10,000, and the 2,000 most variable features were identified using the variance-stabilizing transformation (vst) method. Data were scaled, and principal component analysis (PCA) was applied for dimensionality reduction. Uniform manifold approximation and projection (UMAP) and t-distributed stochastic neighbor embedding (t-SNE) embeddings were computed using the first 20 principal components.

Lineage trajectories were inferred using Monocle3, with cells ordered along pseudotime. Root principal nodes were defined using donor-specific cells (BM D1), and pseudotime values were visualized on UMAP embeddings. Differential gene expression between high and low osteogenic potential MSCs was initially assessed using Seurat’s FindMarkers function. Significant genes were subsequently analyzed using STRINGDB for enriched Gene Ontology (GO) terms and GO term molecular function (top 8).

Pseudobulk expression matrices were generated using AggregateExpression, grouped by donor and osteogenic potential. Differential expression analysis between high and low osteogenic potential groups was performed using DESeq2. Genes with |log₂ fold change| > 1 and adjusted p-value < 0.05 were considered significant. Volcano plots were generated to visualize differential expression, and selected genes were displayed as feature plots on UMAP embeddings.

Cells were clustered using the Louvain algorithm with a resolution of 1.4. This resolution yielded 25 transcriptional clusters, consistent with the degree of heterogeneity previously documented in hBMSC populations^3,47^. Cluster annotation confirmed that the majority of clusters were present across all donors, with donor-specific enrichment concentrated in a restricted subset, confirming that the clustering resolution does not artificially inflate donor-specific transcriptional differences. Combined cluster annotation (clusters 2, 3, and 11) was defined based on UMAP visualization. Differential expression analysis between clusters of interest and all other clusters was performed using Seurat’s FindMarkers, and significant genes were analyzed in STRINGDB to identify enriched pathways. Molecular function GO term images were exported, and dot plots were generated for genes significantly enriched under the GO term “Chondrocyte differentiation” or the “TISSUES” database top-hit chondrocyte expression. The top 2,000 highly variable genes were used to calculate pseudobulk expression profiles for each cluster, averaging the log-normalized RNA counts across all cells within the same cluster. From the differential expression analysis, the 30 most significantly upregulated genes (adjusted p-value < 0.05, positive log fold change) were selected. A heatmap was generated showing these top 30 genes as rows, ordered by decreasing log fold change, and the clusters as columns, hierarchically clustered based on the pseudobulk expression of the 2,000 variable genes.

### Human cell characterization by flow cytometry analysis

As previously reported stromal cells were characterized by their plastic adherence and tri-differentiation potential *in vitro*^7,65^. Moreover, in here hBMSCs were profiled by the expression of consensus stem cell-like surface markers (CD29, CD44, CD49d, CD73, CD90, CD105, CD146, CD34, and CD45) detailed in **Table 1**. Briefly, cells were trypsinized, washed in phosphate buffer-saline (PBS) and resuspended in FACS buffer (1 mM ethylenediaminetetraacetic acid (EDTA) and 2% FBS in PBS) containing each antibody at an optimized concentration. Cells were stained for 30 min at 4 °C in the dark, washed, filtered through a 70 µm filter and analyzed by a BD FACSCanto flow cytometry analyzer. Dead cells were stained by SYTOX Blue dead cell stain and excluded from analysis like doublets. Data analysis was performed using FlowJo 10.0.8 software (TreeStar), and gates were assessed based on the corresponding isotype controls.

### Cell proliferation within TG-PEG hydrogels

hBMSCs from the six different donors were encapsulated in optimized TG-PEG hydrogels (15 µl hydrogels, 1.7% TG-PEG containing MMP-degradable and RGD sites) at a density of 5x10^6^ cells per ml of hydrogel and cultured in MEMα supplemented with 1% P/S, 10% FBS, and 5 ng ml^-1^ of FGF-2. Cell proliferation/metabolic activity was assessed post-encapsulation (time 0), and after 4, 8, 12, and 16 days in culture using the CellTiter-Glo 3D Cell Viability Assay (CTG, Promega) according to the manufacturer’s instructions. Briefly, hydrogels were washed once with serum-free medium and incubated in a 1:1 mixture of culture medium and CellTiter-Glo reagent for 30 min at room temperature under gentle agitation. ATP standards were prepared in parallel to generate a calibration curve. Luminescence was quantified using a microplate reader following transfer to white-walled assay plates.

### Osteogenic differentiation and alkaline phosphatase activity quantification

For osteogenic differentiation in 2D culture, hBMSCs from individual donors were seeded in 24-well plates (22500 cells per well, to match the number of cells in TG-PEG) and maintained in MEMα supplemented with 1% P/S and 10% FBS for 24 h to allow cell attachment. Cells were then cultured for 12 days in osteogenic or control medium. For 3D osteogenic differentiation, hBMSCs were encapsulated in optimized TG-PEG hydrogels (15 µl, 1.7% TG-PEG containing MMP-degradable and RGD sites, with 1.5x10^6^ cells per ml of hydrogel) and cultured in 48-well plates for 12 days. Osteogenic medium consisted of MEMα supplemented with 1% P/S, 10% FBS, 10 mM HEPES, 1 mM sodium pyruvate, 2 mM L-glutamine, 50 µg ml^-1^ L-ascorbic acid, 10 mM β-glycerol phosphate and 100 ng ml^-1^ BMP-2. Control medium lacked BMP-2. Media were exchanged every 4 days. At endpoint, alkaline phosphatase (ALP) activity was quantified. For 3D cultures, hydrogels were washed with PBS and enzymatically degraded using collagenase A (2 mg ml^-1^ in PBS) at 37 °C for 30 min to recover encapsulated cells. Cell pellets were lysed in alkaline lysis buffer (0.56 M 2-amino-2-methyl-1-propanol containing 0.2% Triton X-100, pH 10.0 in H2O) for 30 min on ice. For 2D cultures, cells were lysed directly in the culture wells following PBS washing. Cell lysates were centrifuged at 16000 g for 10 min and the supernatants were collected for ALP quantification. ALP activity was measured colorimetrically using p-nitrophenyl phosphate as substrate. Absorbance was measured at 410 nm using a microplate reader.

### Chondrogenic differentiation and glycosaminoglycan (GAG) quantification

For conventional chondrogenic differentiation, hBMSCs from individual donors were cultured as pellet aggregates (250 000 cells per pellet) in 15 ml conical tubes for 21 days. To assess chondrogenesis in synthetic matrices, hBMSCs were encapsulated in optimized TG-PEG hydrogels (10 µl hydrogels for GAGs, 15 µl hydrogels for histologies, 1.7% TG-PEG containing MMP-degradable and RGD sites, with 20x10^6^ cells per ml of hydrogel) and cultured in 48-well plates for 21 days. Chondrogenic medium consisted of high-glucose DMEM (41965) supplemented with 1% P/S, 1% ITS+ premix, 1 mM sodium pyruvate, 40 µg ml^-1^ L-proline, 50 µg ml^-1^ L-ascorbic acid, 100 nM dexamethasone and 20 ng ml^-1^ TGF-β3. Control medium lacked dexamethasone and TGF-β3. Media were exchanged every 3 days. At endpoint, samples were washed with PBS and either frozen at -80 °C for biochemical analyses or fixed in 4% paraformaldehyde (PFA) for histological evaluation. For histology, hydrogels were embedded in HistoGel (Epredia) prior to paraffin embedding to preserve construct morphology during processing. For glycosaminoglycan (GAG) quantification, samples were digested overnight in proteinase K solution (1 mg ml^-1^ proteinase K in 50mM Tris buffer containing 1mM EDTA, 1mM Iodoacetamide, 10 µg ml^-1^ pepstatin). Sulfated GAG content was quantified using a dimethylmethylene blue (DMMB)-based colorimetric assay with chondroitin sulfate standards and normalized to DNA content measured using the CyQUANT assay (Invitrogen).

### Mass spectrometry analysis of hBMSC-derived ECM

To characterize donor-dependent differences in ECM deposition, hBMSCs with high (BM D1) or low (BM D6) osteogenic potential were encapsulated in optimized TG-PEG hydrogels (25 µl hydrogels, 1.7% TG-PEG containing MMP-degradable and RGD sites, with 20x10^6^ cells per ml of hydrogel). Prior to encapsulation, cells were trypsinized and washed three times with MEMα containing 1% P/S to remove residual serum proteins (**Supplementary Fig. 4a**). Cell suspensions were mixed with TG-PEG precursor solutions and polymerization was initiated by addition of 5 U ml^-1^ of activated human recombinant Factor XIII (rFXIIIa; T080, Zedira). Recombinant FXIIIa was used in place of plasma-derived FXIIIa to avoid contamination from serum albumin present in plasma isolates, which interferes with downstream proteomic analyses^26^ (**Supplementary Fig. 4b**).

Cell-laden hydrogels were cultured in 48-well plates in MEMα supplemented with 1% P/S without FBS for 14 days, with medium changes every 3 days. Acellular TG-PEG hydrogels were included as negative controls. Five independent hydrogel replicates were analyzed per donor condition. At day 14, constructs were washed three times with PBS and enzymatically digested using sequencing-grade trypsin (V511C, Promega; 8 µg ml^-1^ in Tris buffer, 50 µl per gel) for 3 h at 37 °C under agitation (300 rpm). Samples were centrifuged at 300 rcf for 10 min to pellet intact cells, and supernatants containing solubilized ECM proteins and hydrogel components were collected and stored at -80 °C until proteomic processing.

Hydrogel fractions were processed by filter-aided sample preparation (FASP)^73^ as previously described^26^. Briefly, samples were mixed with sodium dodecyl sulfate (SDS) lysis buffer containing dithiothreitol (DTT), heated at 95 °C, clarified by centrifugation, and loaded onto Microcon-30 filters. SDS was removed by sequential washes with urea buffer and NaCl, and free cysteines were alkylated with iodoacetamide. Proteins were digested overnight on-filter with sequencing-grade trypsin in triethylammonium bicarbonate buffer. Peptides were recovered by centrifugation, acidified with trifluoroacetic acid, and desalted using self-packed C18 stage tips. Eluted peptides were dried by vacuum centrifugation and resuspended in 3% acetonitrile and 0.1% formic acid containing iRT peptides.

Peptides were analyzed by LC-MS/MS on a nanoAcquity UPLC M-Class system coupled to an Orbitrap Fusion LUMOS mass spectrometer. Peptides were separated by reversed-phase chromatography on an Acquity UPLC M-Class HSS T3 C18 column (1.8 µm, 75 µm x 250 mm, Waters) maintained at 50 °C using a linear acetonitrile gradient containing 0.1% formic acid. Samples were injected in randomized order, with a standard sample included every four injections for quality control. Full scan MS spectra were acquired from 300 to 2000 m/z at a resolution of 120 000. Precursors selected for MS2 analysis were fragmented by higher-energy collisional dissociation (HCD) using a normalized collision energy of 35 and analyzed in the ion trap. MS acquisition was performed in data-dependent acquisition mode with a 3 s cycle time, excluding precursor ions with unassigned charge states or charges of +1 or greater than +7. Dynamic exclusion was set to 15 s with a mass tolerance of 10 ppm.

To assess residual serum protein contamination, LC-MS/MS datasets were additionally analyzed in Skyline (v19.1.1.248)^74^ using BSA and HSA peptide signatures together with iRT peptide standards. MS1 feature integration was performed using Orbitrap filter settings at 120,000 resolving power, and peptide intensities were log2-transformed for comparative analysis.

Raw files were processed in MaxQuant (v1.6.2.3) using label-free intensity-based absolute quantification (iBAQ)^75^. Trypsin/P was selected as the digestion enzyme, allowing up to two missed cleavages. Carbamidomethylation of cysteine was set as a fixed modification, and oxidation of methionine and deamidation of asparagine and glutamine were set as variable modifications. Protein identification was performed with a protein false discovery rate (FDR) threshold of 0.05, followed by filtering for proteins identified with at least two peptides. Spectra were searched against a database containing human reviewed canonical UniProt entries and known mass spectrometry contaminants.

Statistical analyses were performed in R using SRMService^76^ and limma^72^ packages. Protein intensities were log2-transformed and robust-scaled using median and median absolute deviation (mad) normalization. Differential protein abundance between conditions was assessed using moderated t-tests with Benjamini-Hochberg correction for multiple testing. Proteins identified in acellular hydrogel controls were removed from the dataset. For proteins identified exclusively in one condition, fold changes were estimated using the mean intensity of the lowest 10% detected protein averages in the comparison group. Proteins identified and detected in at least three hydrogels per condition were retained for downstream analysis. ECM-associated proteins were annotated by cross-referencing the resulting protein lists with the Matrisome Project database^77,78^. Volcano plots and heatmaps were generated in R (v3.6.1) using gplots. Functional enrichment analysis of differentially expressed proteins was performed using High Throughput GoMiner^79^ with 1000 randomizations, an FDR threshold of 0.05, and GO category sizes ranging from 5 to 500 genes. GO term-protein matrices were hierarchically clustered using complete linkage and uncentered correlation and visualized with Java TreeView^80^.

### Whole gel immunostaining

To validate proteomics hits, hBMSCs were embedded in optimized TG-PEG hydrogels (15 µl hydrogels, 1.7% TG-PEG containing MMP-degradable and RGD sites, crosslinked with rFXIIIa, and containing 20x10^6^ cells per ml of hydrogel) and cultured in MEMα supplemented with 1% P/S and without FBS for 2 weeks *in vitro*. Hydrogels were subsequently fixed with 4% PFA at room temperature, permeabilized with 0.3% Triton X-100, and blocked in 1% BSA in PBS for 45 minutes at room temperature with shaking at 50 rpm. Primary antibodies (**Table 1**) were then added and incubated overnight at 4°C with shaking at 25 rpm. The following day, gels were thoroughly washed with PBS (3x 1 h washes at 4°C) and incubated overnight at 4°C with secondary antibodies together with phalloidin to F-actin filaments staining and DAPI (1 µg/ml) for nuclear labeling, with shaking at 25 rpm. For collagen type I staining only, samples were incubated with the primary antibody for 5 h in the cell culture incubator prior to fixation. Samples were then washed, fixed with 4% PFA for 20 min, washed three times with PBS, and processed as described above for secondary antibody staining. Finally, gels were thoroughly washed in PBS and imaged using an inverted laser-scanning microscope (Leica TCS SP5). Image analysis and processing were performed using Fiji. Negative controls, consisting of either omission of the primary antibody or isotype-matched controls, were included in all staining experiments.

### Statistical analysis

All data are reported as mean ± standard deviation with individual data points shown as dots. Statistical analyses were performed using GraphPad Prism (version 8.0.0 to 10.0, GraphPad Software). Mean values were compared using an unpaired two-tailed Student’s t-test, or one-way or two-way analysis of variance (ANOVA) followed by Tukey’s or Šídák’s *post hoc* test for multiple comparisons, as specified in each figure legend. Statistical significance was accepted at P < 0.05 and reported as follows: *P < 0.05, **P < 0.01, ***P < 0.001, ****P < 0.0001, NS, not significant. Throughout, N denotes the number of independent biological donors and n denotes the number of technical replicates per condition.

## Data Availability Section

The raw human sequencing data generated in this study have been deposited in the European Genome-Phenome Archive (EGA). Processed single-cell RNA sequencing matrices and Seurat objects have been deposited at Zenodo. All data will be made publicly available upon publication.

All other data supporting the findings of this study are available from the corresponding author upon reasonable request.

## Acknowledgements

This work was supported by the Swiss National Science Foundation (grant CR33I3_153316/1 and CRSII5_186271), and by the Center for Clinical Research, University Hospital Zürich and University of Zürich. We thank Dr. Vincent Milleret (University Hospital Zürich) for project discussions, and Dr. Christopher Millan (University of Zürich) for manuscript editing. We also thank Esther Kleiner (University Hospital Zürich) for technical assistance; Flora Nicholls (University of Zürich) for animal care; Dr. Paolo Cinelli (University of Zürich) for support with microCT; Prof. Marcy Zenobi-Wong and Dr. Nicolas Broguiere (ETH Zürich) for help with rheological measurements; Prof. Franz Weber (University of Zürich) for the synthesis and purification of BMP-2; Dr. Sibylle Pfammatter from the Functional Genomics Center and Dr. Marcel Bühler for their support in proteomics (University of Zürich and ETH Zürich), the Center for Microscopy and Image Analysis for SEM imaging (University of Zürich), Dr. Malgorzata Kisielow and the Flow Cytometry Core Facility (ETH Zürich) for cell sorting assistance, and Julie Russeil (EPF Lausanne) for RNA library preparation.

## Contributions

Q.V.M. designed, performed, and analyzed all experiments. E.A.R. contributed to the proteomics studies including experimental design, sample preparation, and data analysis. L.M., B.M.C, D.A., V.G. and B.D. contributed to the RNA-sequencing experiments including experimental design, sample and library preparation, and data analysis. A.B. and I.M. isolated and expanded primary human cells. M.L. contributed to experimental design and data interpretation. M.E. conceived and supervised the project. Q.V.M. and M.E. wrote the manuscript with input from all authors. All authors discussed the results and approved the final manuscript.

