## Supplemental Figures for "Growth factor-free synthetic matrices reveal intrinsic skeletal progenitor competence"

### Supplementary Information

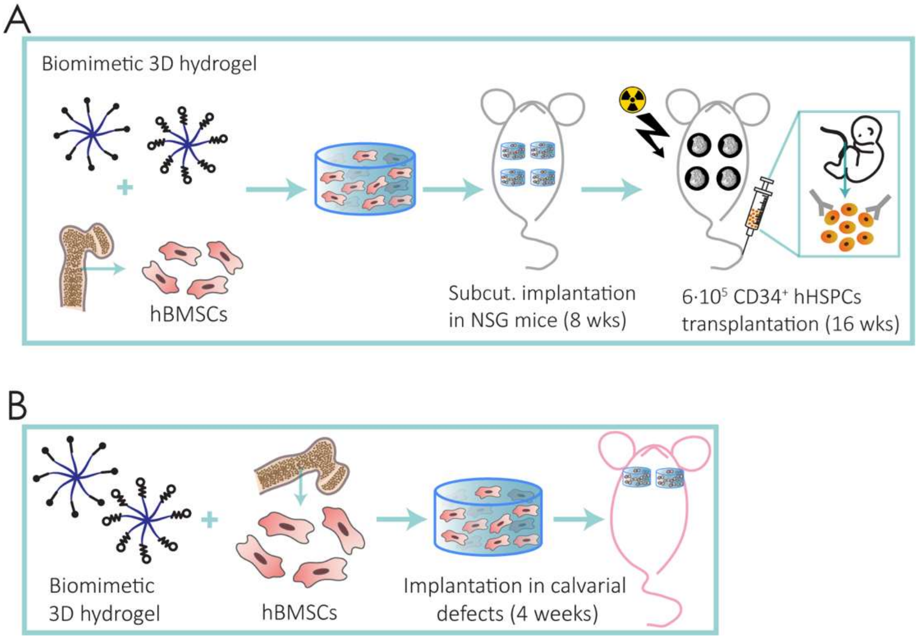


**Figure S1. Functional hematopoietic niche formation and calvarial bone repair by growth-factor-free TG-PEG constructs**

**a,** Schematic of the humanized ossicle model. Bone ossicles were generated by subcutaneous implantation of hBMSC-laden TG-PEG hydrogels in NSG mice for 8 weeks. Human hematopoietic stem and progenitor cells (hHSPCs) were then systemically injected via the tail vein, and hematopoietic engraftment was assessed 16 weeks later.

**b,** Schematic of the critical-sized calvarial defect repair model. A 4 mm diameter full-thickness calvarial defect was surgically created in the parietal bone of nude mice. Optimized TG-PEG hydrogels loaded with hBMSCs (20 x $10^6$ cells ml$^{-1}$) or acellular controls were implanted into the defect site. Bone repair was assessed at 4 weeks post-implantation by microCT volumetric analysis and histological evaluation.

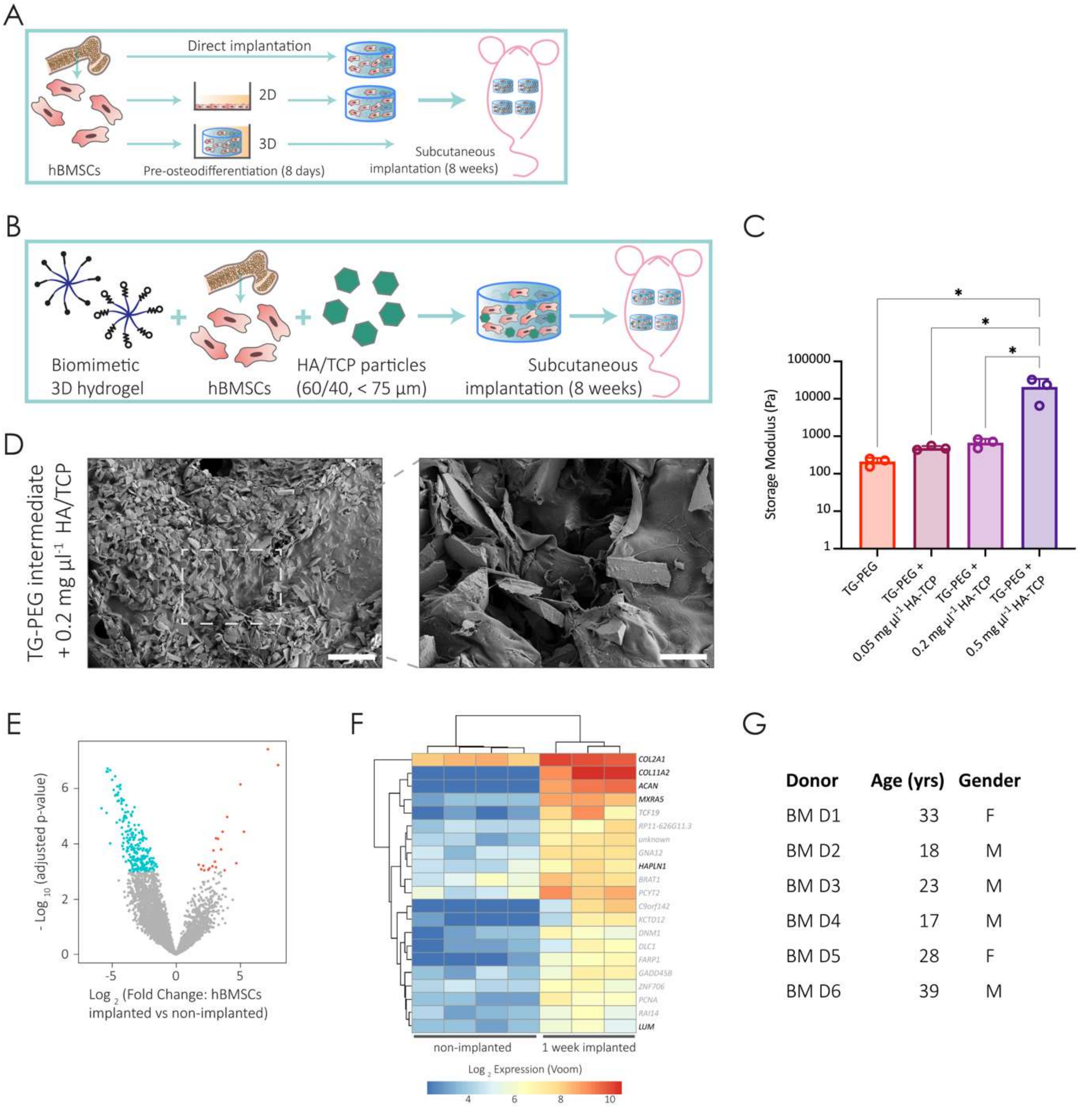


**Figure S2. Controls for osteoinductive priming, osteoconductive particle incorporation, transcriptional profiling, and donor demographics**

**a,** Schematic of the osteogenic pre-differentiation and implantation workflow. hBMSCs were cultured for 8 days in osteogenic medium (MEMα supplemented with 12.5 ng µl$^{-1}$ BMP-2) either in 2D monolayer culture or within 3D TG-PEG hydrogels prior to subcutaneous implantation. Pre-differentiated 2D cells were re-encapsulated in optimized TG-PEG hydrogels immediately before implantation; 3D pre-differentiated constructs were implanted directly. Undifferentiated hBMSCs encapsulated in TG-PEG hydrogels served as positive controls.

**b-d,** Hydroxyapatite/β-tricalcium phosphate (HA/TCP) particle incorporation into TG-PEG hydrogel backbone.
**b,** Schematic of HA/TCP particle incorporation into TG-PEG hydrogels. HA/TCP particles (60/40 ratio, diameter <75 µm) were mixed into TG-PEG precursor solutions at increasing concentrations prior to crosslinking and subcutaneous implantation.
**c,** Effect of HA/TCP particle incorporation on hydrogel mechanical properties. Storage moduli of TG-PEG hydrogels containing increasing amounts of HA/TCP particles, measured by oscillatory rheometry (n = 3 independent hydrogel measurements per condition).

**d**, Scanning electron microscopy of HA/TCP particles incorporated within TG-PEG hydrogels. Representative micrographs confirm homogeneous distribution of HA/TCP particles within the hydrogel matrix. Scale bar left: 400 µm, zoom in right: 80 µm.

**e-f,** Early transcriptional response of hBMSCs to *in vivo* implantation. Human cells were isolated from TG-PEG hydrogels 1 week post-implantation (n = 3 implants) and compared to control non-implanted hBMSCs in TG-PEG scaffolds (n = 4 hydrogels), cells were then analzyed by bulk RNA sequencing.
**e,** Volcano plot showing differentially expressed genes upon *in vivo* implantation (upregulated genes in orange; downregulated genes in blue).
**f,** Heat map of significantly upregulated genes in implanted cells following Voom normalization; extracellular matrix-related genes are highlighted (n = 3 implants for *in vivo*; n = 4 hydrogels for *in vitro* controls).

**g,** Demographic characteristics of the hBMSC donor cohort. Table summarizing age and sex of the six healthy bone marrow donors (BM D1 to BM D6) whose cells were used throughout this study.

Bars represent mean ± s.d.; dots indicate individual hydrogels. One-way ANOVA with Tukey's multiple comparisons test (c). * $P < 0.05$.

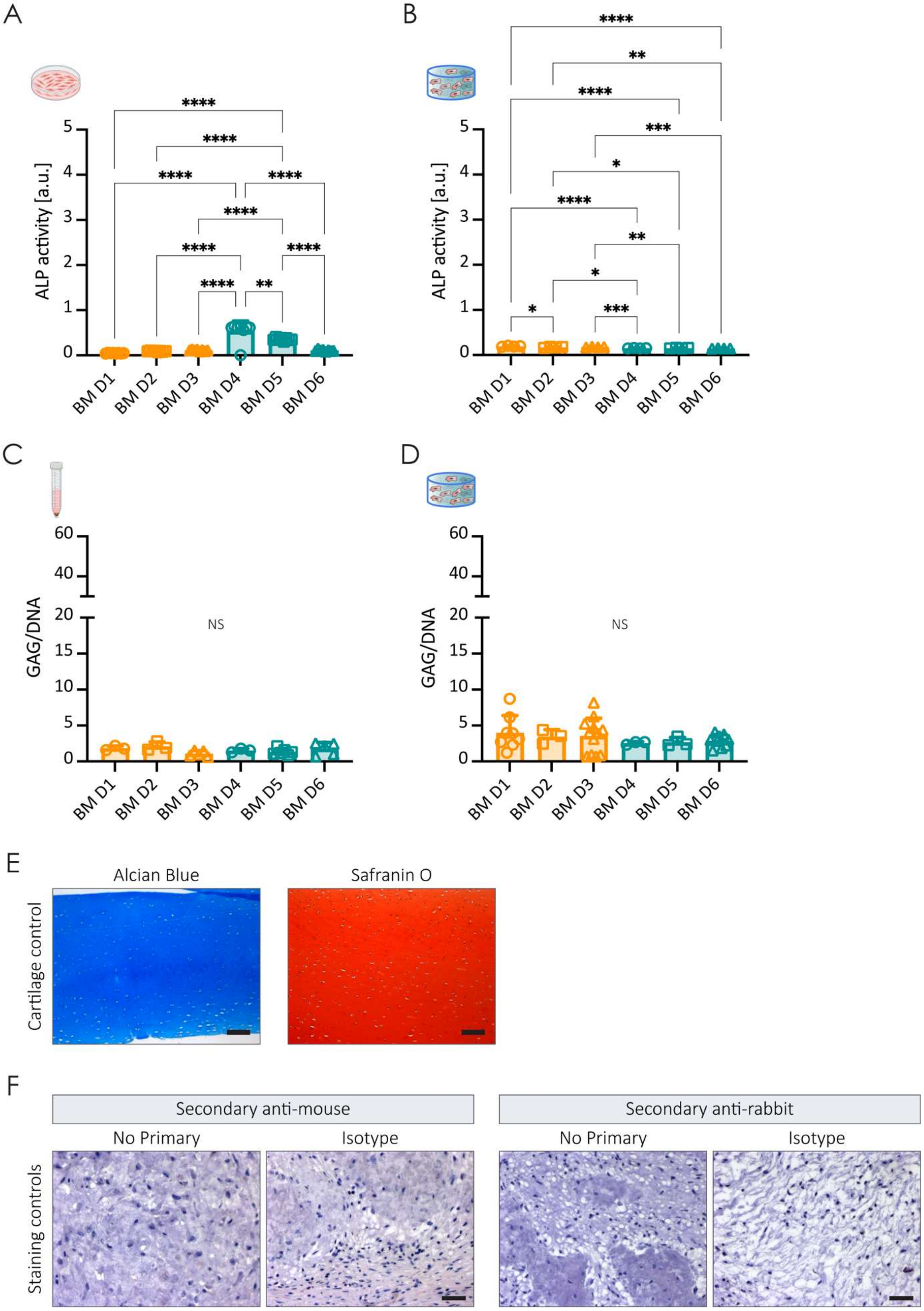


**Figure S3. Basal medium controls confirm specificity of osteogenic and chondrogenic differentiation assays**

**a-d,** hBMSCs from different donors were cultured either in conventional cultures (monolayer or pellets) or encapsulated in 3D TG-PEG hydrogels and maintained under basal medium for 12 days for osteogenic or 21 days for chondrogenic differentiation.

**a,** hBMSCs from all six donors cultured in 2D monolayer under basal medium conditions, quantified by ALP activity. Absence of ALP induction confirms that osteogenic activity measured in **Fig. 4c** is specific to osteogenic medium supplementation (N = 6 donors; n = 8 wells per donor).
**b,** hBMSCs from all six donors encapsulated in 3D TG-PEG hydrogels under basal medium conditions, quantified by ALP activity. Confirms that osteogenic differentiation observed in **Fig. 4d** is medium-dependent (N = 6 donors; n = 4 hydrogels per donor).
**c,** hBMSCs from all six donors in conventional pellet culture under basal medium conditions, quantified by GAG deposition. Confirms that chondrogenic matrix production in **Fig. 4e** is specific to chondrogenic medium supplementation (N = 6 donors; n ≥ 3 pellets per donor).
**d,** hBMSCs from all six donors encapsulated in 3D TG-PEG hydrogels under basal medium conditions, quantified by GAG deposition. Confirms that GAG deposition in **Fig. 4f** is medium-dependent rather than a constitutive property of 3D culture (N = 6 donors; n ≥ 3 hydrogels per donor).

**e**, Positive tissue controls for Alcian blue and Safranin O staining, performed on bovine articular cartilage sections. Alcian blue stains sulfated glycosaminoglycans blue; Safranin O stains proteoglycan-rich matrix red. Scale bars: 100 µm

**f**, Negative controls for immunohistochemical staining. Representative sections processed without primary antibody (no primary control) or with isotype-matched control antibodies in place of primary antibodies, confirming specificity of staining observed in **Fig. 4g** and **Fig. 6**. Nuclei were counterstained with hematoxylin. Scale bars: 50 µm.

Bars represent mean ± s.d.; dots indicate intra-donor replicates (a-d). One-way ANOVA with Tukey's multiple comparisons test (a-d). * $P < 0.05$; ** $P < 0.01$; *** $P < 0.001$; **** $P < 0.0001$; NS, not significant.

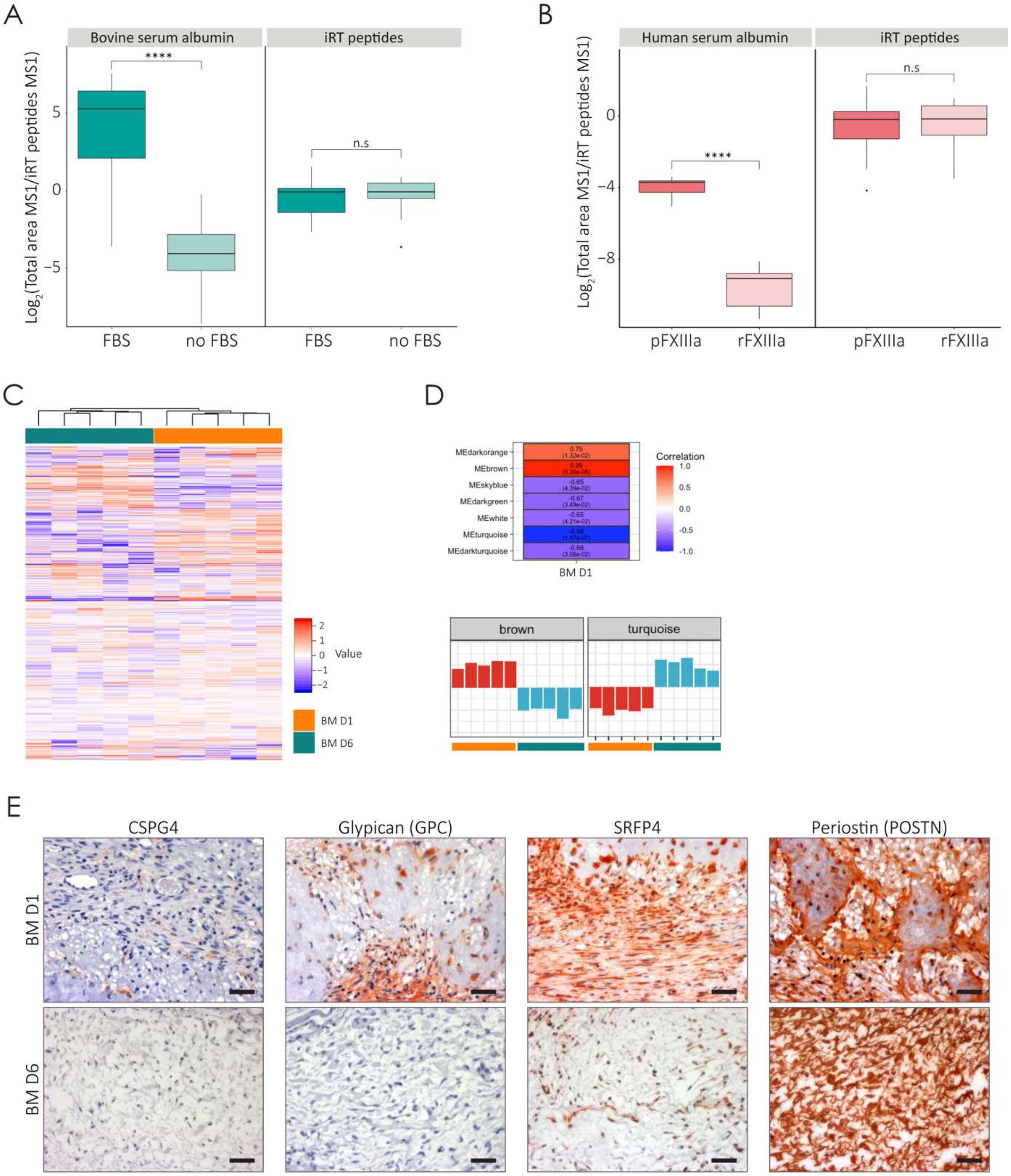


**Figure S4. Optimization and quality controls for proteomic profiling of cell-deposited ECM and *in vivo* validation of donor-specific matrix signatures**

**a-b,** Optimization of TG-PEG hydrogel and culture conditions for downstream proteomics analysis. Serum-free culture conditions and recombinant FXIIIa were validated to eliminate background serum protein contamination that would otherwise interfere with detection of cell-deposited ECM components.

**a, Left:** Quantification of the total MS1 feature area for the best ranked 28 BSA peptides (by spectrum counts) in TG-PEG hydrogels cultured 14 days in medium containing FBS or in FBS-free medium (n = 2 hydrogels). **Right:** Differences in the quantification of iRT peptide intensities among the two samples were non-significant (n = 2 hydrogels). Data are depicted as median (line) surrounded by box plots (25th and 75th percentiles) with whiskers. Lower whisker = smallest observation equal to or greater than lower hinge – 1.5 x IQR (interquartile);

upper whisker = largest observation equal to or less than upper hinge + 1.5 x IQR, (n = 2 hydrogels). Paired two-tailed Student's t-test, **** $p < 0.0001$; n.s.: non-significant.

**b, Left:** Quantification of the total MS1 feature area for the best ranked 8 HSA peptides (by spectrum counts) in TG-PEG hydrogels cross-linked with pFXIIIa or rFXIIIa and immediately digested and processed for LC-MS (n = 2 hydrogels). **Right:** Differences in the quantification of iRT peptide intensities among the two samples were non-significant (n = 2). Data are depicted as median (line) surrounded by box plots (25th and 75th percentiles) with whiskers. Lower whisker = smallest observation equal to or greater than lower hinge – 1.5 x IQR (interquartile); upper whisker = largest observation equal to or less than upper hinge + 1.5 x IQR, (n = 2 hydrogels). Paired two-tailed Student's t-test, **** $p < 0.0001$; n.s.: non-significant.

**c,** Heatmap showing unsupervised protein clustering of all proteomic samples. Samples cluster by donor identity (BM D1 vs BM D6) rather than by replicate, confirming that inter-donor differences in ECM composition exceed technical variability (N = 2 donors; n = 5 hydrogels per donor).

**d,** Weighted gene co-expression network analysis (WGCNA) of cell-deposited ECM proteomes. **Top:** Seven co-expression modules were identified, each associated with a characteristic protein co-expression pattern and color-coded accordingly. **Bottom:** Module eigenprotein expression profiles across individual samples from BM D1 and BM D6 donors. The brown and turquoise modules show the strongest donor-dependent separation and were selected for downstream pathway enrichment analysis shown in **Fig. 5b** (N = 2 donors; n = 5 hydrogels per donor).

**e,** Representative immunohistochemical staining for chondroitin sulfate proteoglycan 4 (CSPG4), glypican-1 (GPC), secreted frizzled-related protein 4 (SFRP4), and periostin (POSTN) in TG-PEG hydrogels containing BM D1 or BM D6 hBMSCs after 2 weeks of subcutaneous implantation *in vivo*. All six proteins identified by *in vitro* proteomics show stronger expression in BM D1 ossicles, confirming that the donor-specific ECM signature established *in vitro* is enacted *in vivo*. Nuclei were counterstained with hematoxylin. Scale bars: 50 μm.
